# Z-Form Stabilization By The Zα Domain Of Adar1p150 Has Subtle Effects On A-To-I Editing

**DOI:** 10.1101/2025.06.02.657529

**Authors:** Parker J. Nichols, Kent A. Riemondy, Jeffrey B. Krall, Jillian Ramos, Raeann Goering, Morkos A. Henen, J. Matthew Taliaferro, Beat Vögeli, Quentin Vicens

## Abstract

The role of Adenosine Deaminase Acting on RNA 1 (ADAR1)’s Z-conformation stabilizing Zα domain in A-to-I editing is unclear. Previous studies on Zα mutations faced limitations, including variable ADAR1p150 expression, differential editing analysis challenges, and unaccounted changes in ADAR1p150 localization. To address these issues, we developed a Cre-lox system in ADAR1p150 KO cells to generate stable cell lines expressing Zα mutant ADAR1p150 constructs. Using total RNA sequencing analyzing editing clusters as a proxy for dsRNAs, we found that Zα mutations slightly decreased overall A-to-I editing, consistent with recent findings. These decreases correlated with mislocalization of ADAR1p150 rather than reduced editing specificity, and practically no statistically significant differentially edited sites were identified between wild-type and Zα mutant ADAR1p150 constructs. These results suggest that Zα’s impact on editing is minor and that phenotypes in Zα mutant mouse models and human patients may arise from editing-independent inhibition of Z-DNA-Binding Protein 1 (ZBP1), rather than changes in RNA editing.

## Background

Life’s long evolutionary history with viruses has left a significant amount of repetitive transposable elements (TEs) in our genomes^1–3^. The vast majority of TEs are found within intergenic regions and are not transcribed^4^, however, there is a significant enrichment of Short Interspersed Nuclear Elements (SINEs), specifically primate-specific Alu elements^5,6^, near or within coding regions^4^. When these TEs are found in 5’ and 3’ Untranslated Regions (UTRs), they have been proposed to mediate intramolecular (between TEs inserted in opposite orientations) or intermolecular (between two RNA molecules) base pairing to form long dsRNAs which can go on to activate dsRNA-sensing pattern recognition receptors (PRRs) such as Melanoma Differentiation Associated Gene 5 (MDA5), a member of the Retinoic Acid-Inducible Gene I (RIG)-Like Receptor (RLR) family of DExD/H-box helicase domain-containing PRRs^7–11^. Other potential sources of inflammatory endogenous dsRNAs include mitochondrial dsRNA (as both strands of human mitochondrial genomes are transcribed^12–14^) as well as dsRNAs formed from convergent or divergent transcription, overlapping genes, and DNA double-stranded breaks^15–18^. MDA5 initiates the interferon (IFN) pathway upon sensing these long dsRNAs (activation requires >300 bps *in vitro*^10,19^ and >∼1000 bps *in vivo*^20^) in the cytoplasm ^7,10,20–22^. To avoid autoimmunity, the cell needs a mechanism to ensure that these endogenous dsRNAs do not aberrantly activate MDA5 when not infected.

To this end, animals encode Adenosine Deaminase Acting on RNA (ADAR) proteins which catalyze the conversion of Adenosine to Inosine (A-to-I) in dsRNAs ^23–25^ (**Figure 1A**). Mutations to human ADAR1 are the basis for interferonopathies, which are diseases characterized by an overactive immune system^26,27^, and the knockout of ADARs is lethal in most cell lines and mice^28–32^. Adenosine nucleotides are converted to inosine through a hydrolytic deamination reaction where water is used to remove adenosine’s amino group in place for a keto group^33,34^ (**Figure 1B**). In addition to its inability to form a strong base pair with uracil, inosine also weakens base stacking interactions within the A-form helix and disrupts minor groove hydration, significantly destabilizing the A-form geometry ^35–37^ and inhibiting MDA5 signaling^10,25,38^ (**Figure 1B**). The vast majority of A-to-I editing sites reside in introns and 3’ UTRs that harbor dsRNA-forming Alu elements^38–44^, which is consistent with the fact that Alu retrotransposons are one of the largest sources of endogenous dsRNAs within the human transcriptome^45^. Editing sites have also been observed to map back to mtRNAs in unstressed cells^46^, as well as highly immunogenic overlapping transcripts derived from adjacent genes transcribed in opposite directions^2^, known as *cis*-natural antisense transcripts (*cis*-NATs)^47^. ADAR1 editing is believed to prevent these endogenous sources of dsRNA from activating MDA5 under normal cellular conditions, particularly Alu elements found within 3’ UTRs^2,10^.

**Figure 1.**
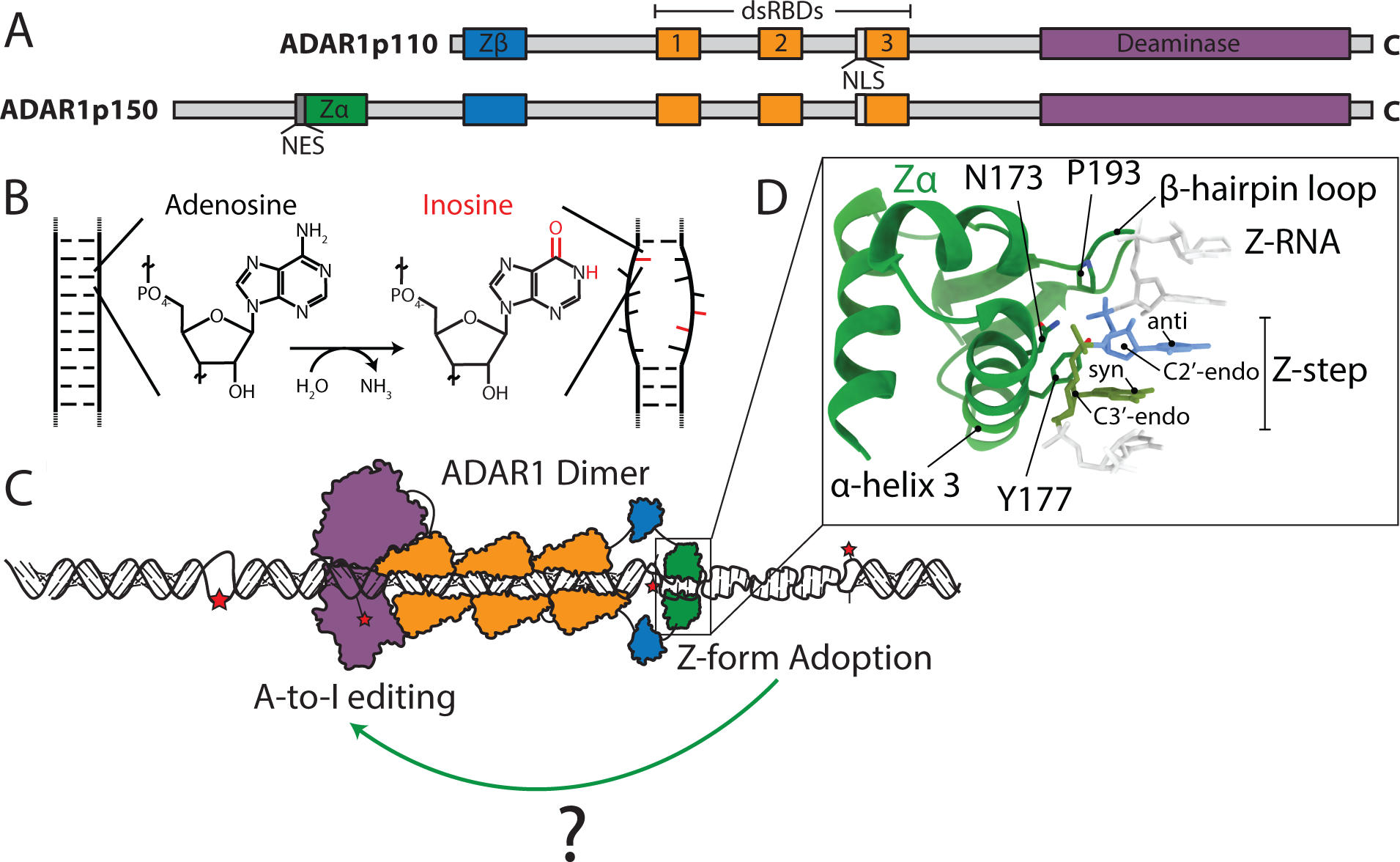
A-to-I Editing by ADAR1 Disrupts A-form Helical Structure and Marks Endogenous dsRNA as Self. (**A**) The domain structure of the short (ADAR1p110) and long (ADAR1p150) isoforms of ADAR1 are shown. ADAR1p110 contains a deaminase domain which is responsible for ADAR1’s catalytic deamination activity, three double-stranded RNA binding domains (dsRBDs) which interact with the A-form structure of dsRNA, a Nuclear Localization Sequence (NLS) and a Zβ domain of unknown function. ADAR1p150 contains the same domain structure but has a ∼300 a.a. N-terminal extension which contains a Nuclear Export Sequence (NES) as well as a Zα domain. (**B**) ADAR1 deaminases adenosine to inosine in dsRNA, which replaces the amino group on the adenosine with a keto group and disrupts A-form helical structure at AU base pairs. (**C**) Cartoon model depicting editing of a dsRNA by ADAR1 and the different domains. (**D**) The Zα domain is able to stabilize the left-handed Z-conformation of dsDNA and dsRNA through key residues which stabilize the unique Z-form geometry.

ADAR1 has two primary isoforms, a shorter, constitutively expressed isoform known as ADAR1p110 which is localized in the nucleus, and a longer, IFN-induced isoform known as ADAR1p150 which is mostly cytoplasmically localized^48–53^ (**Figure 1A**). A-to-I editing is enhanced by the upregulation of the ADAR1p150 isoform during the IFN response^24,49,52^ and is both required and sufficient to prevent aberrant activation of the MDA5/MAVS pathway^54–56^. In addition to ADAR1p150’s cytoplasmic localization, it is also unique in that it harbors an N-terminal Zα domain, which is a winged helix-turn-helix domain that recognizes the higher-energy left-handed conformation of nucleic acids instead of typical B-DNA or A-RNA^57–61^ (**Figure 1A, 1C**).

The Z-conformation is made up of a repeating dinucleotide unit known as a Z-step where the sugar puckers alternate between the C2’- and C3’-*endo* conformations along with the bases between the *anti-* and *syn-*conformations^57,61–64^ (**Figure 1D**). This unique geometry results in a jagged backbone geometry that “zig-zags” along the helical axis and causes the helix to be narrower and more extended compared to the B- and A-conformations. Furthermore, the phosphates between strands are closer together on average, resulting in unfavorable electrostatic repulsion, accounting for a significant amount of the Z-forms instability in the absence of specific buffer conditions or Zα domains^65–69^. Zα domains are able to stabilize the Z-conformation of nucleic acids by making a series of favorable contacts with many of these unique and energetically unfavorable features^59,70^ (**Figure 1D**). In the Zα domain from ADAR1p150, K169 and K170, N173, R174, and T191, all of which are present on the third α-helix of Zα, make a series of charge-charge, water-mediated, and hydrogen bonding interactions with the Z-RNA backbone phosphate residues and the O4’ atoms of the ribose sugars^59,71^. P192 and P193 are important for positioning Zα’s β-hairpin loop and also form van der Waals interactions with Z-DNA/Z-RNA. One of the most critical residues (and often mutated as a means to prevent Z-form stabilization) is Y177 which inserts into the minor groove and forms a CH-π interaction with the nucleotide that adopts *syn* conformation^59,71^.

Several studies have indicated a role for the Zα domain of ADAR1p150 in augmenting A-to-I editing during the IFN response^72–75^. Specifically, point mutations to the Zα domain of ADAR1 which impair its ability to stabilize the Z-form have been shown to result in reduced A-to-I editing of SINE-containing RNAs, and mouse lines harboring these mutations develop spontaneous MAVS-dependent IFN responses^72–75^. This is supported by *in vitro* editing assays which showed that dsRNA substrates containing a Z-form prone sequence are edited at a nearly 40% higher level^76^. How the binding by the Zα domain of ADAR1p150 and Z-form stabilization results in this increase in editing has remained poorly understood.

In contradiction to these findings, other studies have observed no decreases in editing upon mutations to Zα and that the IFN response observed in Zα mutants is editing-independent^77,78^, or that the decrease in editing observed does not explain the observed drastic increase in IFN signaling and cell-death pathways^79^. Indeed, one study demonstrated that the expression of ADAR1p150 constructs containing a variety of Zα mutations, two of which are implicated in Aicardi-Goutières Syndrome (AGS) and Bilateral Striatal Necrosis (BSN)^80,81^, had no obvious effect on ISG signaling through MDA5 activation^2^.

Whether Zα has a role in editing by ADAR1 remains unclear. Previous studies investigating the impact of Zα mutations on A-to-I editing faced some limitations, including variable ADAR1p150 expression levels due to transient transfection, methodological issues in analyzing differential A-to-I editing, and a failure to account for potential changes in ADAR1p150 localization ^72–75^. To address these challenges, we designed a Cre-lox system in ADAR1p150 KO cells to make a collection of stable cell lines expressing different Zα mutant ADAR1p150 constructs. We then carried out total RNA-sequencing of these mutant cell lines and analyzed individual A-to-I editing in addition to employing an analysis strategy similar to one used by the Li lab to identify editing clusters as a proxy for long dsRNAs^2^. We also tracked the localization of the different ADAR150 mutants using fluorescence microscopy. Our results show that mutations to Zα do indeed decrease the overall editing of ADAR1p150, albeit only slightly, in agreement with more recent studies. However, these decreases to editing correlated with mislocalization of ADAR1p150, and we were unable to identify specific sites which were differentially edited between wild-type and the Zα mutants within statistical certainty. These results suggest that the observed decreases are likely not due to lower specificity in the absence of Z-form stabilization and instead are due to changes in ADAR1p150 localization upon mutation of the Zα domain. Our results support the idea that while Zα does appear to have a role in augmenting A-to-I editing, it is likely not to the extent required to cause the phenotypes observed in Zα mutant mice models and in human patients. These phenotypes may be due instead to editing-independent inhibition of Z-DNA-Binding Protein 1 (ZBP1).

## RESULTS

### Rationale for the choice of wild-type and ADAR1 mutants used in this study

To investigate the importance of being able to stabilize dsRNA into the Z-conformation for A-to-I editing by ADAR1p150, we introduced mutations into ADAR1p150’s Zα domain, which are known to impact Zα’s Z-form stabilization properties. In the past, a variety of such mutations have been identified and characterized. The N173S and P193A mutations associated with AGS and BSN map to Zα’s nucleic acid binding helix (α helix three) and β-hairpin loop, respectively, both of which are involved in making interactions with the Z-form helix^59,71^. We have recently analyzed these two mutants by NMR, which revealed that the N173S mutant results in a global disruption of Zα^ADAR1^’s stability and solution dynamics, whereas the P193A mutation, on the other hand, resulted in a 5.4 Å rearrangement of Zα’s β-hairpin loop^82^. Both of these mutations decreased Zα^ADAR1^’s affinity for a canonical Z-forming r(CpG)_3_ duplex by ∼three orders of magnitude and prevented stabilization of a r(CpG)_3_ duplex into the Z-conformation. A double mutation of N173A and Y177A was reported to completely abolish Zα’s ability to stabilize the Z-form while leaving the protein fold intact^83^. This double mutation has been exploited in the *in vivo* setting as a way to prevent Z-form signaling by ADAR1p150^84,85^. We also created a complete Zα deletion as a more blunt method of knocking out Z-form stabilization, in case that the point mutations resulted in subtle effects due to still have an intact domain structure. Lastly, we generated a catalytically dead version of ADAR1p150 (H910Q,E912A, described here^86^), which should not be able to catalyze the conversion of adenosine to inosine and therefore should not result in an increase in editing levels. These mutated ADAR1p150 constructs are illustrated in **Figure 2A**.

**Figure 2.**
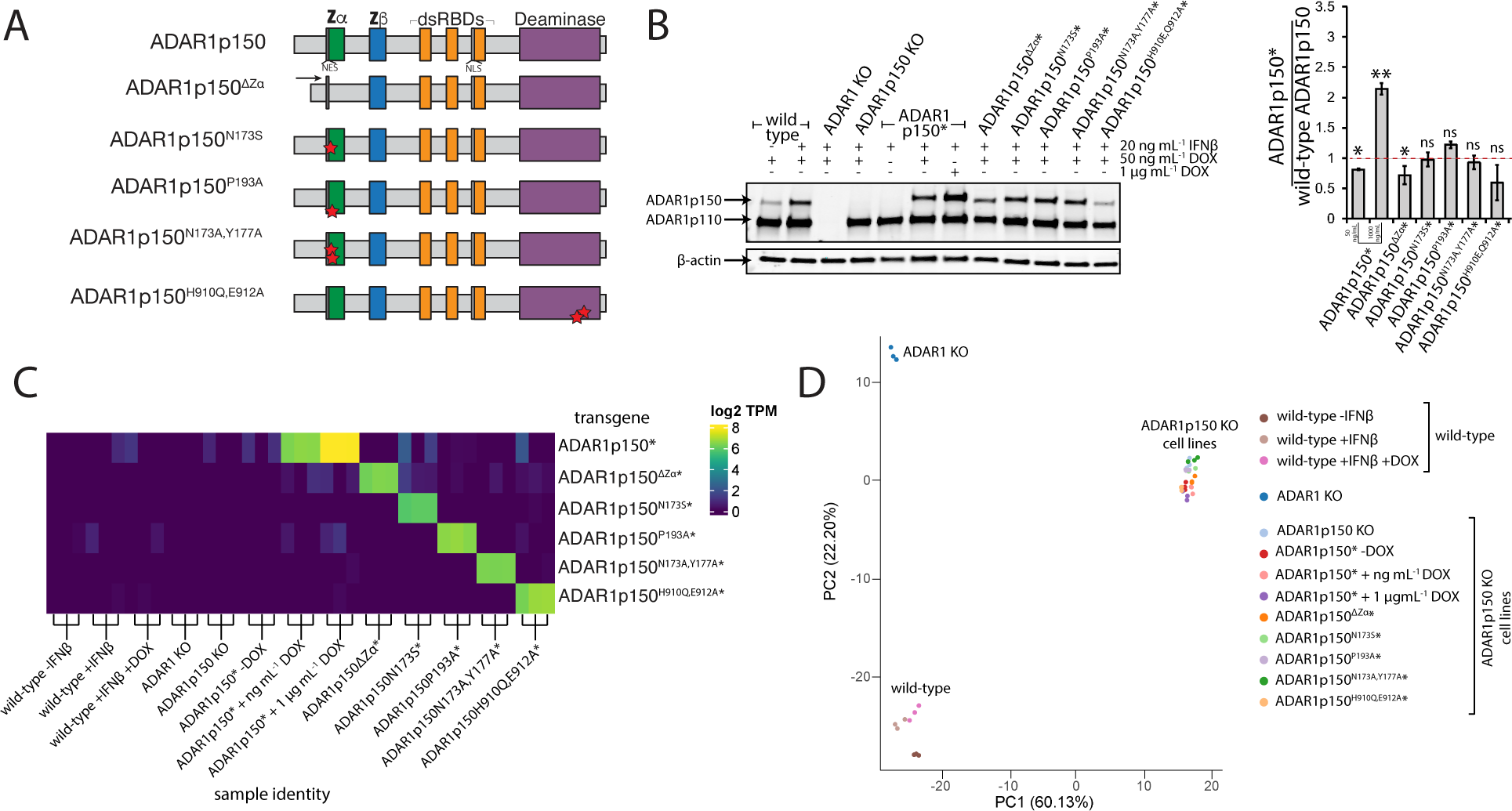
A Cre-Lox System Allows for Tunable ADAR1p150 Mutant Expression in HEK293T Cells. (**A**) Domain architectures and point mutant locations for the different re-integrated ADAR1p150 mutant constructs. Red stars indicated mutation sites. (**B**) Western blot (left) and quantification (right) showing doxycycline-inducible expression levels of the re-integrated ADAR1p150 mutants relative to re-integrated wild-type ADAR1p150. Quantification was from three replicates, all of which are shown in **Supplemental** Figures 5 and 6. (**C**) mRNA expression levels in Transcripts Per Million (TPM, on the *y-axis*) of the re-integrated ADAR1p150 constructs for each cell line from RNA-seq data. For each transgene, the detected expression value is indicated for each sample. (**D**) Principal Component Analysis (PCA) of gene expression of the wild-type HEK293T cells, ADAR1 KO and ADAR1p150 KO HEK293T cells, and the ADAR1p150 re-integrated cell lines.

### A Cre-lox Cassette System in ADAR1p150 KO Cells Allows for the Rapid Creation of ADARp150 Mutant Stable Cell Lines

We used a previously established method to generate stable knock-in cell lines by integrating a lox cassette into the AAVS1 safe harbor locus using CRISPR/Cas9-mediated homologous recombination^87^. The cassette contains a blasticidin resistance gene flanked by loxP and lox2272 sites. Upon Cre-mediated recombination with an incoming donor plasmid, the blasticidin gene is replaced with a puromycin resistance gene, the rtTA3 transactivator, and a doxycycline-inducible expression cassette consisting of Tet operator sequences upstream of a minimal CMV promoter driving the gene of interest (**Supplemental Figure 1A**). This configuration enables stable genomic integration and precise control of transgene expression in response to doxycycline. We applied this approach to ADAR1 KO and ADAR1p150 KO cells which were generously provided by the Rice lab and described in detail elsewhere^88^. After creating the parent cell line with the integrated loxP cassette, creating new cell lines takes ∼2 weeks from transfection to freezing stocks (**Supplemental Figure 1B**). We confirmed successful integration of the loxP cassette into the AAVS1 locus through PCR using primers targeted to the junction between the AAVS1 locus and the integrated cassette, and by integrating EGFP into the locus and confirming that it could be expressed through the addition of doxycycline (**Supplemental Figure 2**).

Employing this approach using the ADAR1p150 KO cells, we generated six stable cell lines expressing wild-type ADAR1p150* (* indicates the construct is re-integrated into the ADAR1p150 KO cells), ADAR1p150^ΔZα^*, ADAR1p150^N173S^*, ADAR1p150^P193A^*, ADAR1p150^N173A,Y177A^*, and ADAR1p150^H910Q,E912A^* under a doxycycline-inducible promoter (**Figure 2A**). We chose to use the ADAR1p150 KO instead of full ADAR1 KO cells to keep the system as close to wild-type as possible and to avoid false positives which could emerge by ADAR1p150 potentially editing sites which would normally be edited by ADAR1p110. Wild-type ADAR1p150 was re-integrated as a positive control to compare the mutants to, whereas the catalytically dead mutant served as a negative control to ensure that expression of ADAR1p150 from a non-endogenous locus did not interfere with the editing levels of ADAR1p110 or ADAR2. To match the expression level of the re-integrated constructs as closely as possible to IFN-treated wild-type HEK293T cells, we incubated the ADAR1p150* cell line with increasing concentrations of doxycycline and analyzed ADAR1p150* expression levels by Western blot, which showed that 50 ng.mL^-1^ doxycycline produced similar protein expression levels to wild-type +IFNβ (**Supplemental Figure 3**). We therefore used this concentration of doxycycline for subsequent experiments.

When using 50 ng mL^-1^ of doxycycline, the expression levels of ADAR1p150^N173S^*, ADAR1p150^P193A^*, ADAR1^N173A,Y177A^*, and ADAR1^H910Q,E912A^* showed no differences within statistical significance (**Figure 2B**, replicates shown in **Supplemental Figure 4 and 5**). ADAR1p150* and ADAR1p150^ΔZα^* levels appeared statistically significant as the expression levels decreased by 19% and 28%, respectively, when compared to wild type levels in HEK293T cells. As expected, treating ADAR1p150* cells with 1 µg mL^-1^ of doxycycline leads to an even higher increase in protein expression levels of about 2.3x compared to wild-type (**Figure 2B**), illustrating a dose-dependence for this expression mechanism. Supporting these findings, we also analyzed the mRNA expression levels of the integrated transgenes from RNA-seq data using Salmon^89^. This analysis showed that the expected constructs were being expressed in the correct cell lines and that the expression levels of the integrated transgenes were largely consistent, except for ADAR1p150^N173S^*, which was lower by approximately 30% compared to ADAR1p150* (**Figure 2C**). The observed differences between protein and mRNA expression levels may be due to the introduced mutations altering the translation efficiency of the constructs, or their stability. Finally, we also checked to ensure that the integration of the various constructs did not cause any significant changes to gene expression profile of the cells. Principal component analysis (PCA) of global gene expression grouped the cell lines according to their genotype –wild-type, ADAR1 KO, or ADAR1p150 KO— demonstrating that insertion of the transgenes into the ADAR1p150 KO cells did not significantly alter their overall transcriptomic profiles (**Figure 2D**).

We further examined the mRNA expression levels of well-known interferon-stimulated genes (ISGs), such as IFNβ1, ISG15, IFIT2, IFIH1, DDX58, and OAS2, to assess the interferon state of the different cell lines. As expected, the ADAR1 KO cells exhibited elevated ISG levels, while all other cell lines showed consistently low and similar ISG expression, indicating that their interferon states are comparable (**Supplemental Figure 6**). ISG15, was elevated in every cell type except for wild-type, consistent with its high sensitivity to IFN stimulation^90^, and has been previously shown to be very sensitive to mutations impacting ADAR1 function^91^. Overall, these findings indicate that re-integration of the mutant ADAR1p150 constructs into the ADAR1p150 KO cells successfully restored mRNA and protein expression levels comparable to wild-type without inducing significant off-target transcriptomic changes. Therefore, we successfully generated a collection of stable ADAR1p150 mutant-expressing cell lines using Cre-lox recombination into ADAR1p150 KO cells, enabling detailed analysis of these mutants.

### Global A-to-I Editing Levels are Minimally Impacted by Zα Mutations

To analyze the impact of mutating Zα on A-to-I editing by ADAR1p150, we incubated the various cell lines with IFNβ for 24 hours. Such conditions mimic the conditions which would normally induce ADAR1p150 expression in wild-type cells^92–95^. In addition, we induced ADAR1p150 expression using 50 ng mL^-1^ of doxycycline and then carried out ribosome-depleted, total RNA-sequencing, with ∼50 million reads per sample. Since inosine preferentially pairs with guanosine, reverse transcription of edited RNAs during library preparation results in the incorporation of cytosine in the complementary strand, which is then read out as guanosine during sequencing ^96^. Consequently, A-to-I editing can be quantified by looking for A-to-G mismatches in the mapped reads compared to a known genome sequence (**Figure 3A**). We first determined A-to-I editing events de novo using a custom pipeline in R to call variants from the mapped RNA-seq reads. However, because 90% of the identified de novo sites from our samples overlapped with known sites from REDIportal (**Supplemental Figure 7**), a comprehensive database of RNA editing sites across various human tissues^97^, we decided to limit our analysis to REDIportal sites to limit false positives.

**Figure 3.**
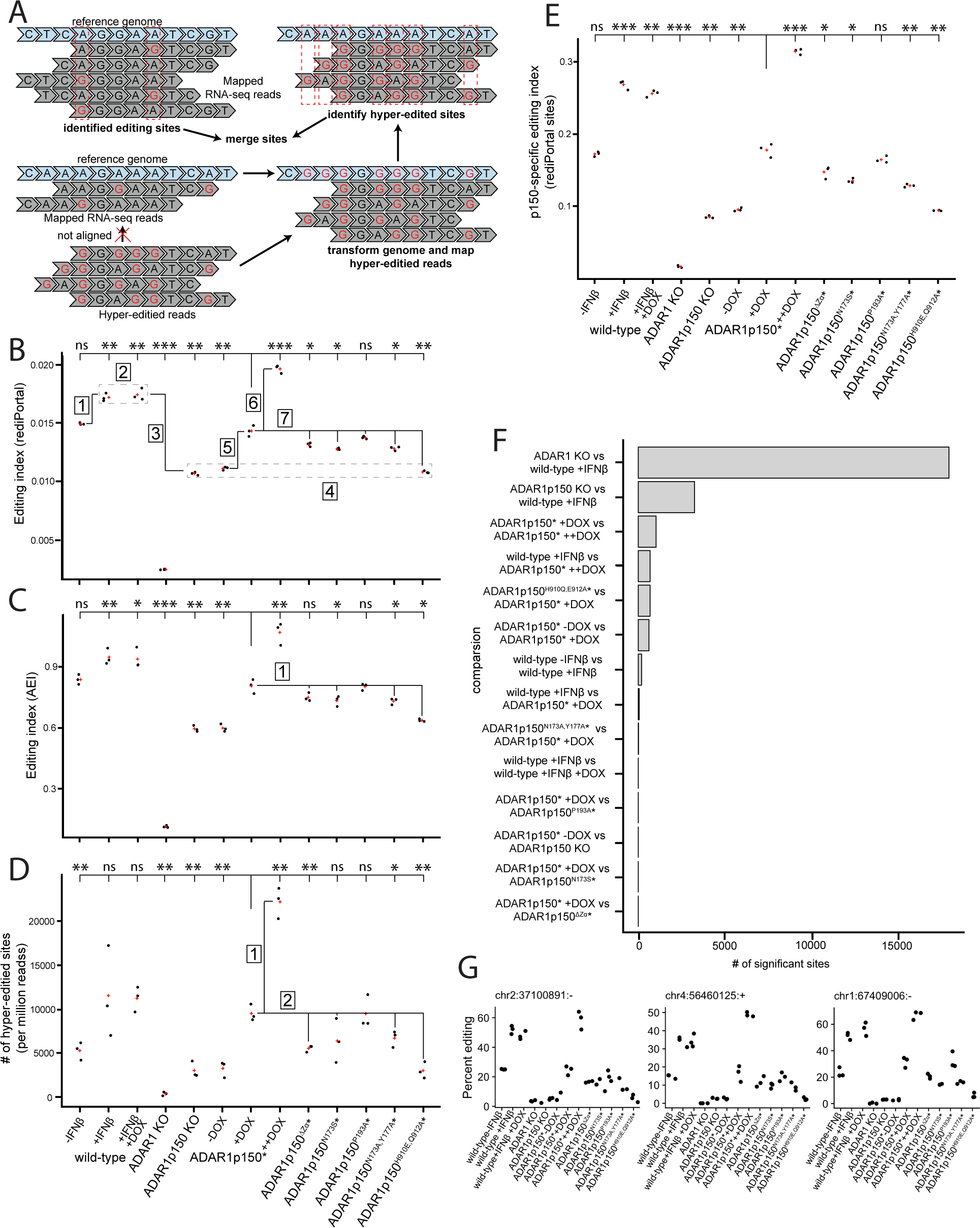
Zα Mutants Result in a Small but Statistically Significant Decrease in A-to-I editing Levels at Individual Sites. (**A**) Illustration of how A-to-I editing events from aligned reads and unaligned reads are identified. Editing indices (the ratio of total A-to-G changes over the total number of adenosines) calculated for all REDIprotal sites (**B**) and Alu elements specifically (**C**, AEI = Alu Editing Index) are shown for the different cell lines. Comparisons referenced to in the main text are marked and numbered. Each black dot represents a replicate and the red cross is the average of three replicates. (**D**) Similar to the editing index, the number of hyper-edited sites per million reads from originally unmapped reads is shown. (**E**) The editing index for ADAR1p150-specific REDIportal sites (by filtering out sites observed in the ADAR1p150 KO cell line) for all cell lines is shown. (**F**) The numbers of differentially edited sites for individual cell line pairs are compared *y-axis* as determined using DESeq2 (**Table 1**). (**G**) Editing frequencies at the top 3 most differentially edited sites between wild-type and ADAR1p150 KO are shown.

**Table 1.**
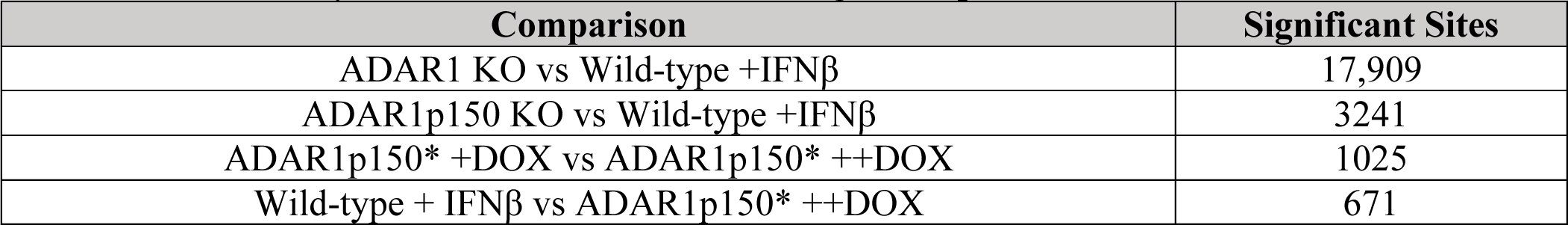
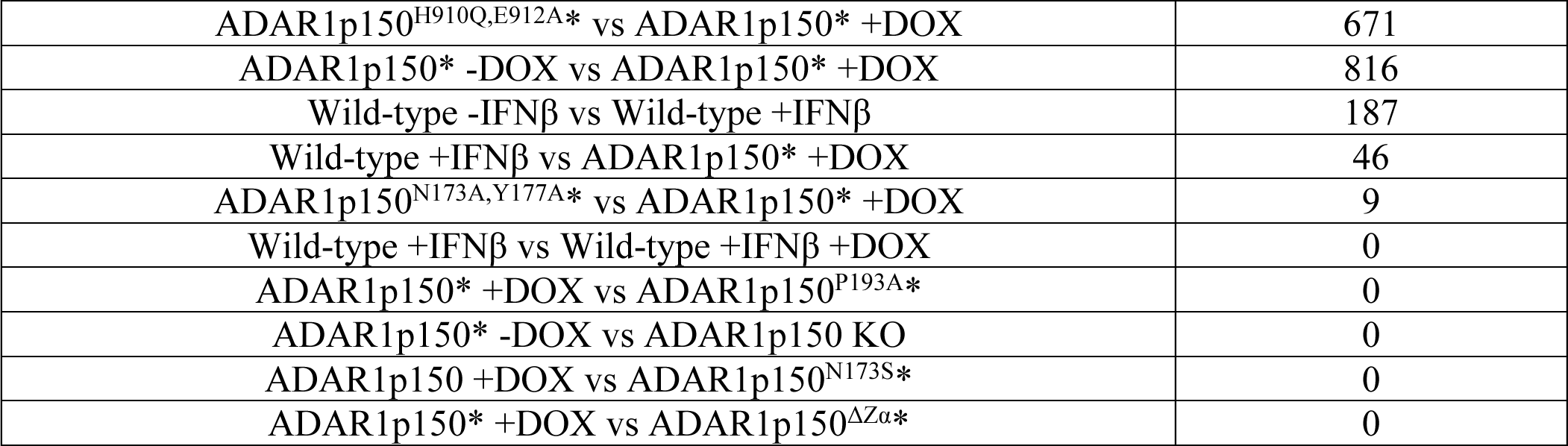
Differentially Edited Sites Determined Using DESeq2.

We determined how the overall editing was impacted by the different mutants by calculating the A-to-I editing index, which is defined as the total number of A-to-G mismatches divided by the total number of possible editing sites (i.e., the sum of unedited and edited adenosines). As expected, treating wild-type cells with IFNβ results in a ∼15% increase in the editing index (**Figure 3B [1]**), consistent with the increased expression level of ADAR1p150, whereas the full ADAR1 KO cells have almost no observed editing and editing in ADAR1p150-specific KO cells is significantly reduced by ∼38% (**Figure 3B [3]**). The addition of doxycycline also does not significantly alter the editing index of wild-type cells or change which sites are edited as determined by DESeq2 analysis (**Figure 3B [2], 3F**), indicating that it should not confound the analysis of the doxycycline-inducible cell lines. Additionally, expression of the catalytically dead ADAR1p150^H910Q,E912A^* results in a nearly identical editing index and edited sites to that of the ADAR1p150 KO cells, indicating that expression from a non-endogenous locus does not alter baseline editing levels and also that the presence of a catalytically dead ADAR1p150 does not act cooperatively with ADAR1p110 to enhance editing (**Figure 3B [4], 3F**). Inducing ADAR1p150* expression in the ADAR1p150 KO cell line rescues editing, however, only to ∼83% of wild-type +IFNβ editing despite having similar protein expression levels to wild-type (**Figure 2B [5]**). This is likely because wild-type ADAR1p150 is expressed at low levels even in the absence of IFNβ, whereas our inducible system only becomes expressed after the addition of doxycycline and therefore does not have enough time to catch up to wild-type editing levels. Indeed, overexpression of ADAR1p150* to more than double that of wild-type levels by treating cells with 1 µg mL^-1^ doxycycline instead of 50 ng mL^-1^ only results in an editing index of ∼114% that of wild-type +IFNβ (**Figure 3B [6]**). Therefore, editing levels appear to be highly sensitive to the concentration and the duration of ADAR1’s expression in the cell.

Because overexpression of the ADAR1p150 mutant constructs might lead to off-target editing, we decided to keep the expression levels of the reintegrated constructs matched to wild-type IFNβ and compare the editing indices of the mutants back to ADAR1p150*, which have similar expression levels (**Figure 2**). Interestingly, all tested mutations except for the ADAR1p150^P193A^* mutant led to statistically significant decreases of the editing index, to levels of ∼91% for ADAR1p150^ΔZα^* and to ∼89% for ADAR1p150^N173S^* and ADAR1p150^N173A,Y177A^* (**Figure 3B [7]**). Since Alu elements are the most significantly edited family of RNAs in the human transcriptome^38–44^, we also looked at how the mutants impacted editing of Alu elements specifically. The overall trend is the same as for all REDIportal sites, with minor decreases to the editing index except that the editing index of the ADAR1p150^ΔZα^* mutant is no longer decreased within statistical significance (**Figure 3C [1]**).

### Analysis of Hyper-Edited Reads Show Similar Minor Decreases in Global Editing Upon Zα Mutations

ADAR1 tends to edit dsRNAs in clusters, a process known as hyper-editing^39,41,42,98–101^. Given that our biophysical characterization of Z-RNA stabilization by the Zα domain of ADAR1 suggested a potential positive feedback loop between A-to-I editing and Zα binding^102^, we wondered whether the Zα domain could have a more pronounced effect on hyper-editing. We recovered and analyzed hyper-edited reads, which initially did not align to the genome due to the high number of mismatches in the sequence, using a previously published pipeline^43^ (**Figure 3A**). Interestingly, the expression level of ADAR1p150 seems to have a more pronounced effect on hyper-edited reads, as treatment with 1 µg mL^-1^ doxycycline, which doubles ADAR1p150 protein levels compared to wild-type +IFNβ, results in a significant increase in the number of hyper-edited sites to ∼192% of wild-type levels, compared to only the ∼114% increase observed for the editing index (**Figure 3D [1]**). ADAR1p150*, when induced with 1 μg/mL doxycycline, is expressed at levels comparable to endogenous ADAR1p110 (**Figure 2B**), yet it results in a ∼6-fold increase in hyper-editing relative to ADAR1p110 alone (**Figure 3D**). This suggests that ADAR1p150 plays a prominent role in hyper-editing, consistent with its established function in suppressing MDA5 activation and in agreement with recent literature^78^. However, this effect may be driven by ADAR1p150’s cytoplasmic localization, where mRNAs spend the majority of their lifespan^103^, and therefore may not be unique to ADAR1p150, a possibility that has been previously suggested ^78^. We also did not test overexpressing ADAR1p110, which may have resulted in similar increases to hyper-editing levels.

Both ADAR1p150^ΔZα^* and ADAR1p150^N173A,Y177A^* show statistically significant reductions in the number of hyper-edited sites, to ∼58% and ∼70% of ADAR1p150* levels, respectively (**Figure 3D [2]**). The high variability among the three replicates of ADAR1p150^N173S^* makes it difficult to determine whether its decrease aligns with the decreases observed in the editing index analysis. Overall, the hyper-editing analysis exhibits greater variability due to the relatively small number of hyper-edited sites (thousands versus millions), which limits the reliability of the results. Therefore, we merged the hyper-edited sites with those from the traditionally aligned reads and focused on p150-specific editing sites by excluding those present in the ADAR1p150 KO condition. The trend closely mirrors that of all sites, with statistically significant albeit minor decreases observed for the ΔZα, N173S, and N173A,Y177A mutations, while the P193A mutation shows no significant difference (**Figure 3E**). Taking these results together, deleting or mutating the Zα of ADAR1p150 so that it can no longer recognize the Z-conformation results in a subtle decrease to overall A-to-I editing while having a more pronounced effect on hyper-editing.

### Zα Mutations Do Not Decrease Editing at Specific Sites

Since global editing patterns of the Zα mutants were lower compared to wild-type ADAR1p150*, we attempted to identify the sites which are differentially edited using DESeq2, which models count data using a negative binomial distribution and applies multiple testing correction to control for false discovery rate^104^. As expected, comparing the ADAR1 KO and ADAR1p150 KO cells to the wild-type +IFNβ revealed the highest number of differential sites with 17,909 and 3241 sites, respectively (**Figure 3F**). Supporting the modest 15% increase in the editing index when treating wild-type cells with IFNβ, only 187 differentially edited sites were identified when comparing wildtype with and without IFNβ, suggesting that the baseline level of ADAR1p150 is sufficient to edit most p150-specific sites. Additionally, only 46 differentially edited sites were found between wild-type +IFNβ and cells expressing re-integrated wild-type ADAR1p150*, indicating that despite the ∼17% lower editing index in ADAR1p150* cells, most sites are still edited at levels similar to wild-type or have editing frequencies which are highly variable compared to the overall decrease to editing (**Figure 3F**). While 618 differentially edited sites were observed between ADAR1p150* cells treated with and without doxycycline, analyzing the Zα mutants revealed that only the ADAR1p150^N173A,Y177A^* cell line had differential editing sites with statistical certainty, and even then, only a modest total of 9 sites (**Figure 3F**, **Table 1**). This number jumps to 1025 differentially edited sites when comparing ADAR1p150* cells treated with 50 ng mL^-1^ and 1 µg mL^-1^ doxycycline, again highlighting that the concentration of ADAR1p150 has a significant effect on editing levels, as seen with the editing index. Thus, the modest decrease in overall editing observed with the Zα mutations is not confined to specific sites but is distributed across all potential editing sites, causing any potential sequence-specific signatures to be obscured by the variability in editing at individual sites. This is further supported by the observation that the same global trends seen in the editing index are mirrored in the editing frequencies of the top 12 most differentially edited sites between wild-type and ADAR1p150 KO, indicating that the effect is widespread rather than driven by reduced editing at a specific subset of sites (**Supplemental Figure 8 and Figure 3G**). Taken together, these findings demonstrate that while the Zα mutations in ADAR1p150 lead to subtle, globally distributed reductions in A-to-I editing, the overall editing landscape remains largely intact, with no significant loss of editing at specific sites. In other words, the Zα domain of ADAR1p150 is not primarily responsible for targeting A-to-I editing to specific, Z-form adopting sequence contexts.

### Editing Clusters are also Minimally Impacted by Zα Mutations

Recently, Sun et al. analyzed the editing status of entire dsRNAs by identifying clusters of editing sites rather than looking at individual sites^2^ (**Figure 4A**). The benefit of this approach is that it allows for differences in overall editing activity that cannot be ascribed to single (and likely low frequency) sites to be determined. An editing cluster is defined as having at least five editing sites, with adjacent sites separated by less than 120 nucleotides, meaning that analyzing clusters is more likely to capture true dsRNAs^2^. To investigate if potential effects of the Zα mutants were being missed by analyzing individual editing sites, we employed this approach on the merged data set containing traditionally-detected and detected hyper-edited sites and identified 77,012 unique clusters across all samples, the vast majority of which were found in repeats (98%), specifically Alu elements (96%), in good agreement with Sun et al.^2^. Editing indices determined at the cluster level mirrored those from analyzing individual sites (**Figure 4B**), albeit the magnitude of the changes was greater for all samples. Treating wild-type cells with IFNβ led to a 50% increase in the cluster editing index (**Figure 4B [1]**) compared to 15% when analyzing sites at an individual level. Expression of ADAR1p150* in the ADAR1p150 KO rescues to ∼74% of wild-type levels instead of 83% (**Figure 3B [2]**). The Zα mutations also show larger decreases compared to ADAR1p150* corresponding to ∼79% for ADAR1p150^ΔZα^*, 73% for ADAR1p150^N173S^*, and 69% for ADAR1p150^N173A,Y177A^* (**Figure 4B [3]**). ADAR1p150^P193A^* again was not significantly different.

**Figure 4.**
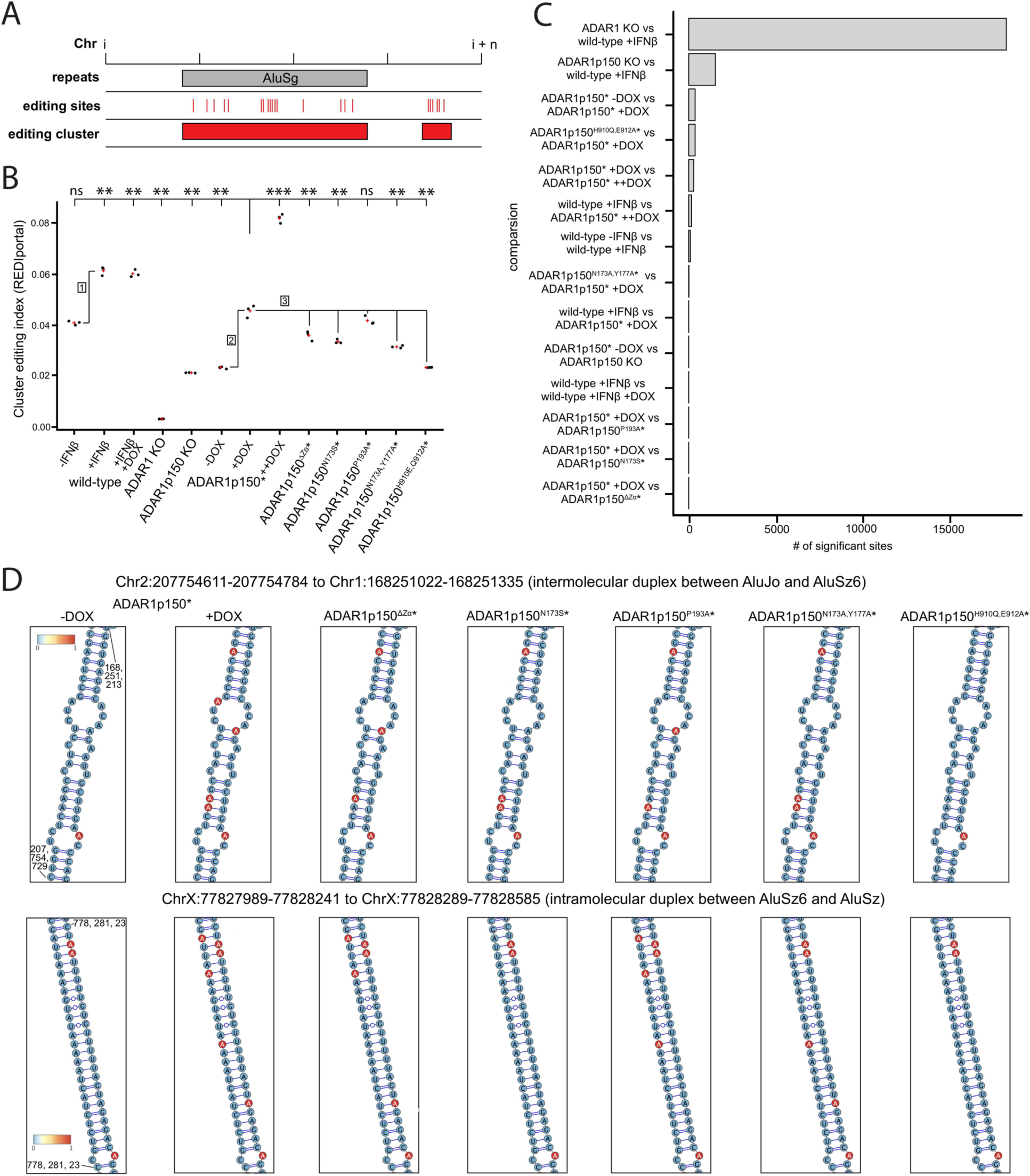
Zα Mutants Result in a Minimal Decrease to Editing at A-to-I Editing Clusters. (**A**) Illustration of how editing clusters are identified, figure adapted from^2^. (**B**) The cluster editing index, calculated by taking the number of A-to-G changes over the total number of adenosines per identified cluster, then averaged for all clusters. Comparisons which are mentioned in the main text are marked and numbered. Each black dot represents a replicate and the red cross is the average of three replicates. (**C**) The number of differentially edited clusters for individual cell line pairs are compared on the *y-axis* as determined using DESeq2 (**Table 2**). (**D**) Zoomed in fragments of Alu foldbacks predicted by the CRSSANT software package^105^ that were also ADAR1p150-dependent (ie, they have differential editing when comparing the ADAR1p150* plus and minus doxycycline conditions). Editing sites were only plotted on the predicted dsRNAs if they had an average editing frequency of > 0.05 averaged over the 3 replicates.

**Table 2.** Differentially Edited Clusters Determined Using DESeq2.

| Comparison | Significant Clusters |
| --- | --- |
| ADAR1 KO vs Wild-type +IFNβ | 18,226 |
| ADAR1p150 KO vs Wild-type +IFNβ | 1534 |
| ADAR1p150* -DOX vs ADAR1p150* +DOX | 376 |
| ADAR1p150<sup>H910Q,E912A*</sup> vs ADAR1p150* +DOX | 373 |
| ADAR1p150* +DOX vs ADAR1p150* ++DOX | 290 |
| Wild-type + IFNβ vs ADAR1p150* ++DOX | 156 |
| Wild-type -IFNβ vs Wild-type +IFNβ | 97 |
| ADAR1p150<sup>N173A,Y177A*</sup> vs ADAR1p150* +DOX | 15 |
| Wild-type +IFNβ vs ADAR1p150* +DOX | 11 |
| ADAR1p150* -DOX vs ADAR1p150 KO | 1 |
| Wild-type +IFNβ vs Wild-type +IFNβ +DOX | 0 |
| ADAR1p150* +DOX vs ADAR1p150<sup>P193A*</sup> | 0 |
| ADAR1p150 +DOX vs ADAR1p150<sup>N173S*</sup> | 0 |
| ADAR1p150* +DOX vs ADAR1p150<sup>ΔZa*</sup> | 0 |

To assess whether any editing clusters were differentially edited as a result of the Zα mutations, we again employed DESeq2, this time comparing cluster editing frequencies instead of editing at individual sites. Consistent with our analysis of individual sites, we observed almost no significantly differentially edited clusters across the mutants (**Figure 4C**, **Table 2**). The N173A,Y177A mutant exhibited only 11 differentially edited clusters, while all other mutants showed none. The lack of significant differentially edited clusters indicates that there is likely not a small subset of dsRNAs missed in the analysis which are specifically impacted by the Zα mutations. The other observed trends also remain similar; comparing wild-type treated with and without IFNβ leads to 97 differentially edited clusters, there are only 11 differentially edited clusters between wild-type +IFNβ and ADAR1p150* +doxycycline, and there are 376 differentially edited clusters between ADAR1p150* treated with and without doxycycline (**Figure 4C**, **Table 2**). Interestingly, comparing ADAR1p150* treated with +50 ng mL^-1^ or 1 µg mL^-1^ doxycycline does not result in an increase of differentially edited clusters in contrast to the observed increase in number of sites. This indicates that the increase in sites due to the higher concentration of ADAR1p150* are potentially distributed more randomly and therefore not captured within clusters. Therefore, similar to our analysis of individual sites, the tested Zα mutants led to an overall decrease of A-to-I editing at identified clusters, however, these decreases were of low magnitude and appeared to be distributed randomly enough such that no or very little differentially edited sites are detected. Illustrating this, we plotted two of the top differentially-edited clusters from Alu foldbacks when comparing ADAR1p150* treated with and without doxycycline onto their secondary structures (**Figure 4D**, plotted on predicted Alu foldbacks from crosslink-ligation data^105^). The editing patterns between the different Zα mutants generally involve the same sites, whereas the catalytically dead mutant is identical to when ADAR1p150* is not expressed (**Figure 4D**).

### Mutations to Asparagine 173 of the Zα Domain of ADAR1 Moderately Alters Localization

Recent work has shown that the unique functions ascribed to ADAR1p150 compared to ADAR1p110 are primarily due to ADAR1p150’s cytoplasmic localization, as both ADAR1p110 and ADAR2 can rescue ADAR1p150 loss by making them cytoplasmic^2,78^. Additionally, the Zα domain of ADAR1 has been implicated in the localization of ADAR1p150 to SGs, an observation which is dependent upon having functional Z-form stabilizing residues^106,107^. Therefore, we wanted to investigate whether any of our selected mutants had altered localizations which could potentially explain the differences observed in overall editing efficiencies.

We employed CRISPR/Cas9 to endogenously tag G3BP1, a DNA helicase which localizes to and promotes SG assembly^108^, with Enhanced Green Fluorescent Protein (EGFP) in wild-type, ADAR1 KO, and ADAR1p150 KO cells (prior to integration of the different ADAR1p150 mutants) so that co-localization of the different ADAR1p150 mutant constructs with SGs could be tracked. We confirmed successful tagging of G3BP1 with EGFP through PCR, western blot, and fluorescence microscopy to confirm that the EGFP-G3BP1still localized to SGs (**Supplemental Figure 9**).

We then treated the cells the same way as for RNA-sequencing (incubation with IFNβ and doxycycline for 24 hours) but also incubated cells with sodium arsenite to induce SGs, and then carried out immunofluorescence with an ADAR1p150-specific antibody. Wild-type cells showed a significant increase in cytoplasmic fluorescence after incubation with IFNβ for 24 hrs, in line with the increased protein levels observed by Western blot, whereas the ADAR1 KO and ADAR1p150 KO cells had little detectable fluorescence (**Figure 5A**). We also observed co-localization between the ADAR1p150 antibody and SGs as marked by G3BP1 upon their induction by sodium arsenite in the wild-type cells but not the ADAR1 and ADAR1p150 KO cells, consistent with ADAR1p150’s known localization to SGs. These results suggest that the ADAR1p150 antibody is indeed specific for ADAR1p150 and sufficient for use in immunofluorescence assays. We also observed for ADAR1 KO and ADAR1p150 KO cells, incubation with IFNβ led to the formation of SGs even in the absence of sodium arsenite, as was previously observed^109^, suggesting that ADAR1p150 is needed to prevent SG formation in the absence of viral infection or oxidative stress.

**Figure 5.**
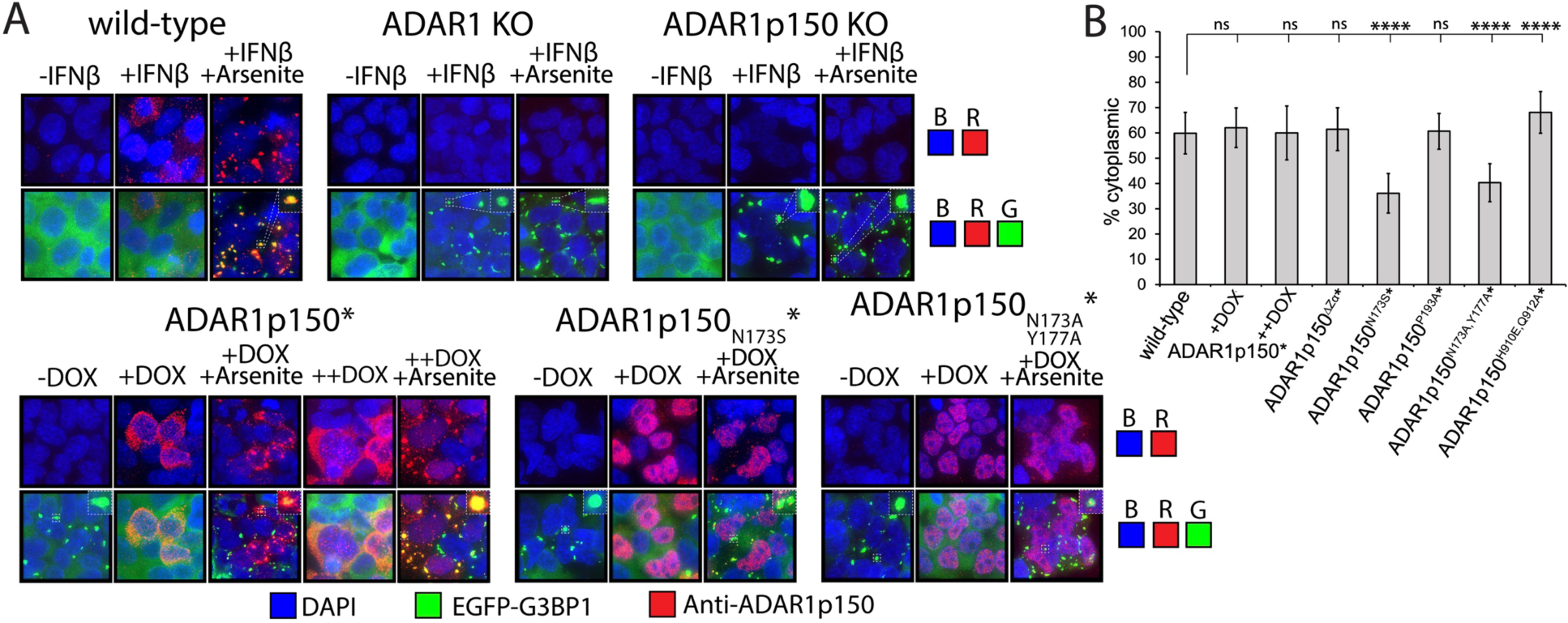
Mutations to N173 of the Zα Domain of ADAR1 Result in Nuclear Retention. (**A**) Immunofluorescence images of the wild-type, ADAR1 KO, ADAR1p150 KO, re-integrated ADAR1p150 (ADAR1p150*), the N173S mutant (ADAR1p150^N173S^*), and the N173A,Y177A (ADAR1p150^N173A,Y177A^*) double mutant cell lines. The red signal is of an ADAR1p150-specific rabbit monoclonal antibody visualized using an Alexa Fluor 594 nm secondary antibody. The green signal is from G3BP1 which was endogenously tagged with EGFP, and the blue signal is DAPI. (**B**) The percentage of the red signal intensity (corresponding to the anti-ADAR1p150 antibody) in the cytoplasm versus the nucleus of the cell.

There is little observable fluorescence from the ADAR1p150 antibody in the ADAR1p150* cells when not treated with doxycycline, however, upon incubation with 50 ng mL^-1^ doxycycline for 24 hrs, we observed a significant increase in cytoplasmic fluorescence consistent with the doxycycline-dependent expression of ADAR1p150* which also co-localized to SGs (**Figure 5A**). Therefore, the re-integrated ADAR1p150* construct behaves nearly identical to wild type, as expected. A doxycycline-dependent emergence in fluorescence was also observed for the other mutants tested as well (**Supplemental Figure 10**). When comparing the different mutants to ADAR1p150*, all constructs looked similar (mostly cytoplasmic and localized to SGs), except for the ADAR1p150^N173S^* and ADAR1p150^N173A,Y177A^* constructs, which localized primarily to the nucleus (**Figure 5A, Supplemental Figure 10**). To quantify this, we measured the relative fluorescence intensity of the different ADAR1p150 constructs inside and outside the nucleus, determining the percentage of each construct localized to the nucleus versus the cytoplasm. The ADAR1p150*, ADAR1p150^ΔZα^*, and ADAR1p150^P193A^* constructs showed approximately 60% cytoplasmic localization, which was not significantly different from wild-type. In contrast, the ADAR1p150^N173S^* and ADAR1p150^N173A,Y177A^* constructs exhibited around 40% cytoplasmic localization, significantly deviating from wild-type (**Figure 5B**). To confirm that this observation was not a potential artifact of antibody-based detection, we also re-integrated and analyzed the localization of mCherry-tagged constructs (ADAR1p150-mCherry*, ADAR1p150^N173S^-mCherry*, and ADAR1p150^N173A,Y177A^-mCherry*), which confirmed the altered localization of the N173S and N17A,Y177A mutants (**Supplemental Figure 11**).

This altered localization was also correlated to less signal intensity observed within arsenite-induced SGs (**Figure 5A**). Since the ADAR1p150^ΔZα^* construct still co-localizes to SGs similar to wild-type, the reduced signal of the N173S and N173A,Y177A mutants in SGs is likely due to lower levels of cytoplasmic ADAR1p150 rather than a loss of Zα-dependent localization to SGs. However, further experiments restoring the localization of the mutants and testing their SG co-localization would be necessary to confirm this. Interestingly, the only other mutant that showed statistically significant differences in its cytoplasmic/nuclear localization compared to the wild type was the catalytically dead H910Q,E912A double mutant, which displayed a slight increase in its cytoplasmic localization compared to wild type (**Figure 5B**). Since this mutant cannot edit dsRNA, we speculate that its altered localization is due to a higher amount of paired dsRNA in the cytoplasm which would promote ADAR1p150 binding.

We wondered why the N173S and N173A,Y177A mutants were the only mutants to have altered cytoplasmic/nuclear localization. Upon inspection of the crystal structure of the Zα domain of ADAR1 in complex with Z-RNA, we noticed that N173 makes a water-mediated hydrogen bond with W195 which forms the hydrophobic core of the domain (**Figure 6A**) and our previous characterization of the N173S mutant by Nuclear Magnetic Resonance suggested that this mutation was destabilizing to the domain’s fold and dynamics^82^. Since ADAR1p150’s nuclear export sequence (NES) partially overlaps with the Zα domain (residues 125-137, with Zα beginning at residue 133), we wondered whether mutations that disrupt Zα stability might also impair nuclear export. Indeed, the partial mis-localization observed in N173S and N173A,Y177A mutants looked very similar to a previous study in which the NES of ADAR1p150 was deleted^110^. To test this, we moved the NES approximately 100 residues upstream of the Zα domain in the N173S construct (NES moved) to see if this would restore proper localization (**Figure 6B**). However, the NES moved mutant exhibited nearly the same disrupted localization as the N173S mutant, thereby having no rescue effect (**Figure 6C, 6D**). This suggests that the altered localization of the N173S and N173A,Y177A mutations is either intrinsic to disruption of the Zα domain and not related to disruption of the nearby NES, or that movement of the NES from its wild-type position results in a similar disruption of nuclear export.

**Figure 6.**
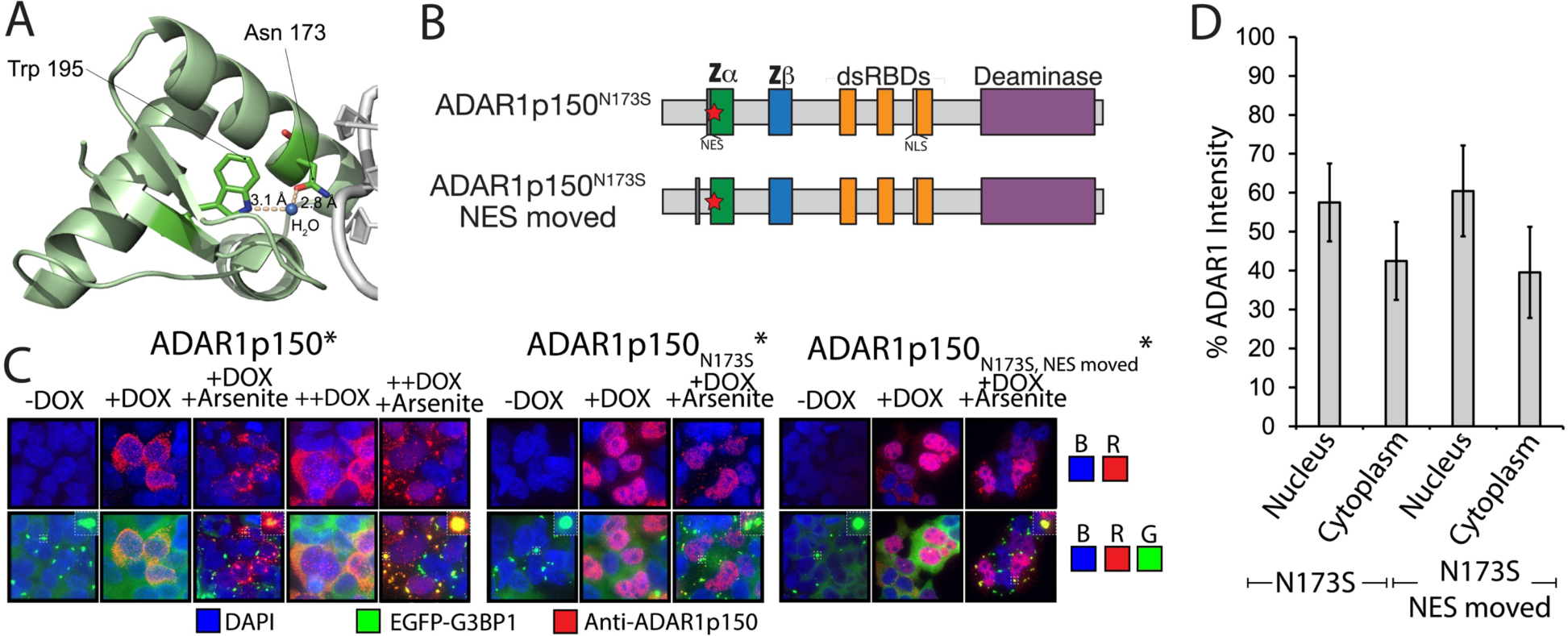
Mis-localization of N173 Mutants is Not Rescued by Relocation of the Nuclear Export Sequence. (**A**) The structure of the Zα domain of ADAR1 bound to Z-RNA is shown (PDB: 2GXB) highlighting the water-mediated hydrogen bond between N173 and W195. (**B**) Domain structures of the ADAR1p150^N173S^ and ADAR1p150^N173S^ NES moved constructs. (**C**) Immunofluorescence images of the re-integrated wild-type ADAR1p150*, ADAR1p150^N173S^, and ADAR1p150^N173S^* NES moved cell lines. (**D**) percentage of Alexa Fluor 594 signal intensity (corresponding to anti-ADAR1p150 antibody staining) measured in the cytoplasm versus the nucleus of the cell for the ADAR1p150^N173S^ and ADAR1p150^N173S^* NES moved cell lines.

Therefore, the N173S and N173A,Y177A mutations moderately alter the localizations of ADAR1p150, going from mostly cytoplasmic to mostly nuclear. Interestingly, the N173S and N173A,Y177A double mutant constructs also showed the largest decreases to editing indices from the RNA-seq analysis (**Figure 3B-E**, **Figure 4B**). Therefore, these decreases are likely due to ADAR1p150’s altered localization in these cell lines instead of altered substrate preferences due to an inability to recognize the Z-conformation, in line with recent work showing that the localization of ADAR1 proteins is primarily responsible for their editing activites^2,78^.

## DISCUSSION

In this study, we investigated the role of the Zα domain of ADAR1 on A-to-I editing by ADAR1p150. A current hypothesis is that Zα is able to enhance editing specificity and selectivity, through stabilizing the left-handed conformation of dsRNA regions^57–61^. We created several stable HEK293T cell lines expressing a collection of ADAR1p150 mutants, under an inducible promoter and analyzed their A-to-I editing abilities by RNA-seq. Specifically, we tested a Zα deletion mutant (ΔZα), two mutants associated with AGS and BSN^80,81^ (N173S and P193A), and a double substitution of N173 and Y177 to alanine which are two of the most critical residues in Z-form stabilization^59,71^. We also analyzed a catalytically dead deaminase construct as a negative control (H910Q,E912A)^86^ (**Figure 2A**). These mutants were chosen because they represent the majority of known mutations that affect Z-form stabilization and analyzing them collectively within a single study allows for direct comparisons between the mutants to be made.

Analysis of our RNA-seq data showed that all of the Zα mutants except for P193A had minor (less than 10%), but statistically significant decreases to global A-to-I editing levels (**Figure 3B-E**), however, we were unable to identify specific sites which were impacted within statistical certainty (**Figure 3F**). This is because the mutations led to a global reduction in editing (**Figure 3G**), but the magnitude of the decreases at individual sites was not greater than the inherent variability in editing frequencies across replicates, making it difficult to pinpoint specific sites affected by the mutations. This reflects the stochastic and variable nature of A-to-I editing within repeat elements^111^. While our re-integrated ADAR1p150* construct restored ∼83% of wild-type editing levels, it remains possible that a subset of editing sites was not recovered within the remaining 17%. However, given the widespread editing observed and the lack of consistent site-specific losses across replicates, we believe this is unlikely to have substantially influenced our conclusions.

We also analyzed editing in dsRNA clusters, which is more sensitive to the editing status of entire dsRNAs, rather than at individual sites^2^ (**Figure 4A**). Analysis of editing clusters mirrored that of individual sites (**Figure 4B**), again showing that while overall editing for all Zα mutants except for P193A was attenuated, no specific sites which had decreased editing could be identified (**Figure 4C)**. We also found that the N173S and N173A,Y177A mutant constructs, which had the largest observed decreases to overall editing (∼89% of wild-type levels) had perturbed localization compared to wild-type (**Figure 5**). Therefore, as observed previously, the localization of ADAR1 proteins appears to have the largest effect on A-to-I editing^2,78^. We observed mislocalization of the N173S and N173A,Y177A mutants in our experiments, likely accounting for their reduced editing activity compared to the P193A mutation, which was properly localized and had no observable effect on editing indices (**Figure 3-5**). In contrast, the decreased editing seen in the ΔZα mutant, despite its correct localization, can potentially be attributed to the complete loss of the Zα domain, and consequently, the loss of an RNA-binding domain—regardless of whether the RNA adopts an A- or Z-conformation. Notably, we previously demonstrated that ADAR1’s Zα domain can recognize A-form helices with low micromolar affinity, suggesting that Zα can bind both A- and Z-form RNAs, though it does so with weaker affinity for A-form RNA^112^.

The other largest contributing factor to overall editing frequencies was the concentration of ADAR1p150 (**Figure 3 and 4**), although our data suggest that these new editing sites were distributed more randomly as judged by the lack in increase of editing clusters. While we attempted to match the expression levels of the mutant constructs to those of wild-type, it is possible that subtle differences in expression could be responsible for the observed editing decreases based on these findings. Indeed, while not statistically significant, minor deviations in the ADAR1p150 expression levels observed by Western blot may explain the small observed differences (or lack thereof) in A-to-I editing levels (**Figure 2B**). Based on the mean, P193A had the highest expression level of the mutants compared to wild-type, which could have masked the subtle impact of mutations on editing. The strong dependence of RNA editing on ADAR1p150 concentration may also explain why mutations in the Zα domain of ADAR1p150 must be paired with alleles that result in the loss of ADAR1p150 expression to give rise to the phenotypes associated with AGS and BSN^80,81^. In conclusion, while we observed minor, but global decreases to A-to-I editing due to the Zα mutations (except for the P193A mutation), Zα does not appear to be important for editing specific, Z-form-dependent sequences within the context of our experimental setup.

Our findings are in agreement with recent studies which also show little to no significant changes in A-to-I editing due to the P193A and W195A mutations^77–79^, and that a functional Zα was not important in inhibiting MDA5-dependent increases to ISG-expression in HEK293T cells^2^. However, other studies have reported increases in IFN signatures, overall reductions in editing, and the presence of differentially edited sites following the mutation of N175 and Y179A (equivalent to N173 and Y177 in humans) or W197A (equivalent to W195 in humans) in mouse ADAR1^72,73^, as well as from the mutation of N173 and Y177A in human ADAR1 in HEK293T cells^74^. Our findings that mutations involving N173 result in altered localization of ADAR1p150 and therefore a larger decrease in overall A-to-I editing may explain some of these findings.

Alternatively, it is possible that the observed increases to IFN signatures are instead due to ZBP1 activation, as is discussed below. Indeed, the observed decreases in A-to-I editing in two of the previously mentioned studies were relatively minor^73,74^, and on-par with the overall decreases to editing observed in our study. These studies also observed differentially edited sites whereas we uncovered very little to none. We employed DESeq2, which is a more conservative method for differential editing analysis, as it carefully models RNA-seq count data, accounts for biological variability, and includes multiple testing correction, reducing the likelihood of false positives^104^. While this approach may detect fewer sites, it provides greater confidence that the identified differences are biologically relevant. Indeed, a follow-up study performed high-coverage re-sequencing of the same N173A,Y177A double mutant ADAR1p150 HEK293T cell line that had originally shown a significant number of differentially edited sites^74^. In contrast to the initial findings, this study observed relatively few differentially edited sites and instead reported a global decrease in editing across all sites^113^. Although some Zα mutations led to an approximately 10% reduction in overall editing, we did not directly assess whether this modest decrease affects MDA5 signaling. Future studies could explore whether such changes have any functional consequences.

Another possibility is that the experimental conditions in our study, as well as those in the other referenced experiments^2,77–79^, did not capture the specific contexts in which Z-form-dependent editing becomes relevant. Recent work showed that RNA extracted from cells undergoing oxidative stress was an extremely potent activator of ZBP1 when transfected back into healthy cells^114^. Since bulky adducts to the C8 carbon of purines can induce the Z-conformation through stereo-electronic effects^115–117^, it is possible that oxidative damage of dsRNA during conditions of oxidative stress (such as 8-oxoguanine (o^8^G)^118^), could create the conditions needed for Z-RNA adoption and result in Zα-dependent changes to ADAR1p150’s editing preferences.

### An Editing-Independent Role for the Zα Domain of ADAR1 in Regulating the Innate Immune Response

If having a functional Zα domain has a minor effect on A-to-I editing by ADAR1p150, why do mutations to Zα often result in significant inflammatory signatures and cell death in mice models and human patients? Potentially explaining this apparent contradiction, recent work has begun to define an editing-independent role of the Zα domain of ADAR1p150 in inhibiting signaling by Z-DNA Binding Protein 1 (ZBP1), a PRR which harbors two Zα domains of its own and initiates multiple forms of cell death upon sensing Z-form nucleic acids in the cytoplasm^119^. ZBP1 senses ligands via its two Zα domains (**Figure 7A**), which enables its RIP Homotypic Interaction Motifs (RHIMs) to oligomerize with each other and with RHIMs from downstream effector proteins, including Receptor-Interacting Protein Kinase 3 (RIPK3). This oligomerization activates Mixed Lineage Kinase Domain-Like Protein (MLKL), thereby initiating multiple forms of cell death^120^. If ADAR1 was solely responsible for editing-dependent inhibition of MDA5, then presumably knocking out both proteins should rescue the negative phenotypes associated with ADAR1 KO. However, it has been demonstrated that ADAR1 and MDA5 double knockout mice typically die within the first few weeks after birth^55,75,121^, but this phenotype can be rescued by mutating key Z-form stabilizing residues in ZBP1 (N46A, Y50A in Zα1, and N122A, Y126A in Zα2)^113,122–124^. Similarly, when the Zα domain of ADAR1 is mutated (N175A, Y179A) in a wild-type ZBP1 background, mice die within two days, but expressing both mutant ADAR1 and ZBP1 prevents mice from dying^113,123^. Moreover, it has been shown that ADAR1p150 and ZBP1 co-immunoprecipitated from cells treated with IFNβ, but not when the Zα domains of ZBP1 are deleted or mutated^122,125^, suggesting that the Zα domains of ADAR1p150 and ZBP1 are interacting with the same RNA substrates and likely competing for binding. This competition also appears to be independent of A-to-I editing, as a catalytically dead ADAR1p150 was able to suppress ZBP1 activation^124^. These lines of evidence suggest that the Zα domains of ADAR1 and ZBP1 directly compete for nucleic acid ligands to regulate the cell’s inflammatory state (**Figure 7B**, model 2), as opposed to augmenting A-to-I editing through altering ADAR1’s affinity or specificity for dsRNA targets (**Figure 7B**, model 1).

**Figure 7.**
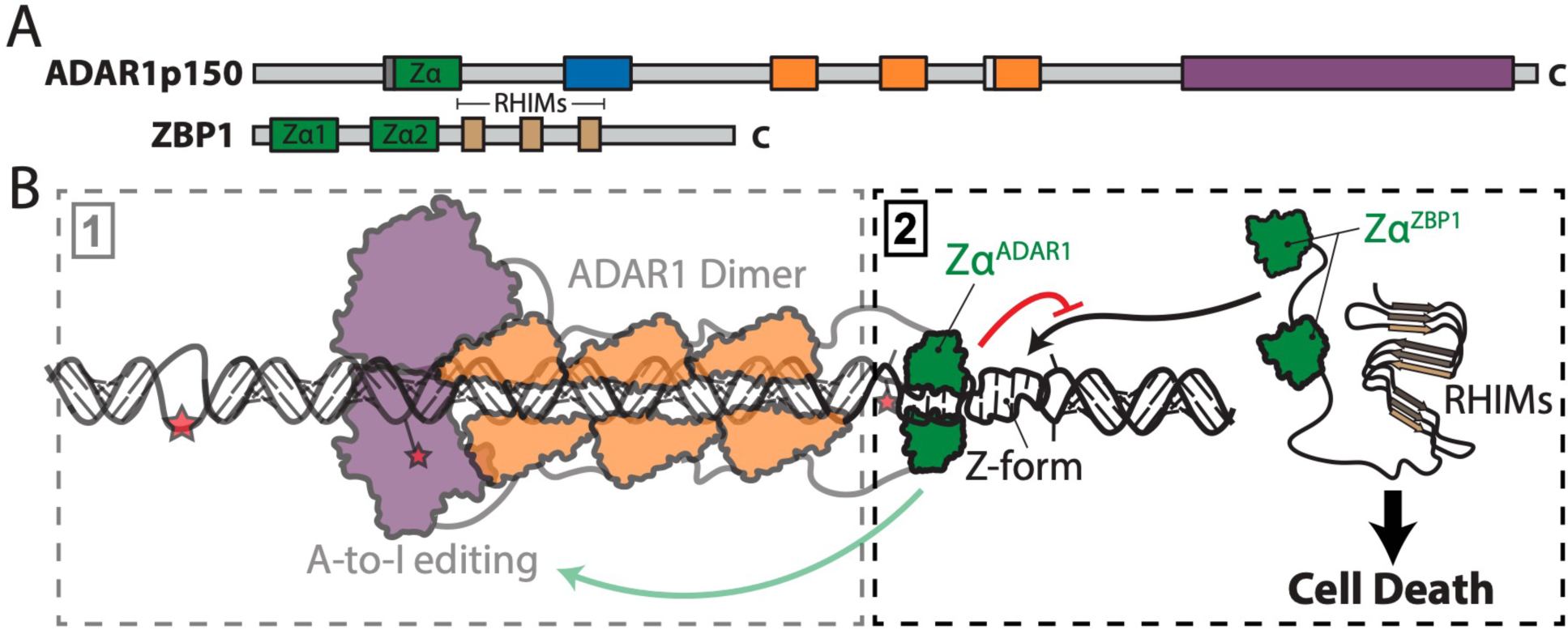
Model for the Role of the Zα Domain of ADAR1 in Regulating Innate Immunity. (**A**) Domain structures of ADAR1p150 and ZBP1 are shown. (**B**) A cartoon depiction of the two potential models showing the effect of the Zα domain on ADAR1 function. In model 1, the Zα domain augments A-to-I editing broadly in a sequence-independent manner. In model 2, the Zα domain of ADAR1 competes with ZBP1 for binding to Z-form substrates in an editing independent manner, thereby inhibiting cell death pathways.

ZBP1 deficiency or mutation to the Zα domains of ZBP1 has also been demonstrated to reduce ISG signatures in ADAR1 mutant cell lines and mice, suggesting that ZBP1 may directly regulate the expression of specific ISGs and inflammatory genes or indirectly contribute to their increased expression through the release of damage-associated molecular patterns from dying cells^113,122,123^. Therefore, it is possible that many of the previously observed changes in ISG signatures upon mutation of the Zα domain of ADAR1 which were prescribed to an impaired ability to edit dsRNA and inhibit MDA5 were instead due to ZBP1 activation^72–75^.

It is also interesting to note that because A-to-I editing is highly specific to dsRNA, it can serve as a readout to confirm whether an RNA is double-stranded and whether ADAR1 was bound to it^126,127^. Therefore, the lack of obvious changes in RNA editing due to mutation of the Zα domain of ADAR1 suggests that it binds many, if not all of the same dsRNA targets regardless of the presence of a functional Zα domain, ie, that the Zα domain is brought along to dsRNA targets rather than playing a role in target selection. Indeed, while the affinity of Zα^ADAR1^ for a pre-stabilized Z-conformation nucleic acid is in the low nanomolar range (4 nM^128^), its affinity for non-Z-form duplexes is three orders of magnitude weaker (4 μM^129^) and exhibits relatively slow conversion kinetics^130^. Therefore, the Zα domain would likely be initially outcompeted for substrate targetting by the low nM binding affinity attributed to the tandem dsRBDs of ADARs^131–133^. This would suggest that the Z-RNA targets of ADAR1p150 and ZBP1 are not unique sequence contexts, but are instead adopted within many of the dsRNA targets which have already been identified, including 3’ UTRs that harbor dsRNA-forming Alu elements^38–44^, mtRNAs^46^, and cis-NATs^2^. We recently determined that sequence constraints of Z-form adoption are looser than originally believed and that unless chemically modified to stabilize the Z-form, Z-conformations are actively induced by Zα domains^102,134^. Therefore, Z-form adoption through the Zα domain of ADAR1 could be potentially thought as acting by a more passive mechanism, where the domain is brought to typical inflammatory dsRNA targets of ADAR1 and then stabilizes Z-conformation stretches within that dsRNA. This could help ADAR1 compete with ZBP1 for Z-RNA binding and potentially help shield the dsRNA from other dsRNA sensors.

We believe it also worthy to note that while the Zα domain of ADAR1 has been shown to be able to stabilize both Z-DNA and Z-RNA^135,136^, recently identified Zα domains from giant viruses are incapable of stabilizing Z-RNA unless they are modified, for example at the C8 position of guanine^137^, which allows the RNA to adopt the Z-conformation even in the absence of Zα or other stabilizing conditions^102^. Therefore, the Z-form adopting targets which are competed for by ADAR1p150 and ZBP1 may be heavily modified dsRNAs that emerge under conditions of extreme cell stress^114^ or may not even be dsRNA but dsDNA instead. Standard RNA-seq based approaches would miss these potentially informative targets.

## METHODS

### Cell Lines and Cell Line Maintenance

The wild-type HEK293T cells were purchased from Millipore-Sigma, and the ADAR1 KO and ADAR1p150-specific KO cell lines were kindly provided by the Rice lab (Rockefeller University)^38^. All subsequent stable cell lines expressing the different ADAR1p150 constructs were generated through Cre-dependent recombination into the loxP cassette within the AAVS1 safe harbor, described below.

All cells were routinely propagated in DMEM/high-glucose medium (HyClone) supplemented with 10% FBS (“characterized” grade; HyClone), 1 mM sodium pyruvate (Invitrogen), and 1× penicillin-streptomycin (100 units mL^-1^ penicillin, 100 μg mL^-1^ streptomycin; Invitrogen). For passaging, adherent cells were detached using 1× trypsin-EDTA (Invitrogen), incubated at 37°C for 3-5 minutes until cells were visibly rounded and lifted from the surface, followed by neutralization with media and dilution into new flasks at the required split ratio or seeding density, depending on the experimental needs, and incubated at 37°C with 5% CO_2_ to maintain optimal growth conditions. Cells were counted using an Invitrogen Countess automated cell counter using Trypan blue following the manufacturer’s instructions.

Cell lines generated in this study were stored in 90% FBS, 10% DMSO upon completing selection, and were rethawed and cultured for no more than two weeks prior to experiments. Frozen cell stocks were generated by gently centrifuging the cell suspension after resuspension in media post trypsinization (800 RPM for 5 min), aspirating the media, resuspending the pellet in 90% FBS, 10% DMSO, and freezing the cells at −80°C overnight. The following morning, frozen vials containing cells were transferred to liquid nitrogen.

### Creation of EGFP-G3BP1 Cell Lines

The G3BP1 donor plasmid for CRISPR/Cas9 of EGFP into the endogenous locus of G3BP1 contained 800 bp homology arms on each side of the insert sequence. The PAM sequence was mutated to prevent recutting of the edited DNA. This plasmid was synthesized by GenScript and was confirmed by full plasmid sequencing (Plasmidsaurus). gRNAs 200 and 209 (**Supplemental Table 1**), targeted towards the first exon of G3BP1 near the start codon and predicted using CHOPCHOP^138^, were cloned into the px330 vector^139^ (acquired from Addgene) after restriction digestion of px330 with BbsI, phosphorylation of the gRNA oligos with T4 PNK, and ligation into px330 using T4 DNA ligase.

ADAR1 KO and ADAR1p150 KO cells were split into 6-well plates (2 mL of media per well) at a seeding density of 300,000 cells per well and incubated for 12-18 hours before being co-transfected either with px330 and the G3BP1 donor plasmid (1 µg each) or with px330 alone as a negative control using Lipofectamine LTX reagent (Invivogen). 10 µL of lipofectamine reagent was combined with 150 µL of OPTI-MEM (ThermoFisher) along with 2.5 µL of plus reagent, plasmid DNA, and 150 µL of OPTI-MEM. The two mixtures were vortexed, then allowed to incubate for 10 minutes at room temperature before combining, vortexing again, and allowing to incubate for another 10 minutes, before adding to cells. The cells were maintained and expanded for one week to get ∼20 million total cells after which GFP-positive cells were sorted into 96-well plates using a XPD100 cell sorter. Single colonies were allowed to recover for ∼1 week before expanding. The negative control cells showed no GFP signal. Correct integration of EGFP into the G3BP1 locus was confirmed by PCR amplifying the junction between the native G3BP1 locus and the inserted EGFP sequence (primers pairs shown in **Supplemental Table 1**), western blotting (the same procedure as for western blotting of ADAR1 was used, see below) using a rabbit anti-G3BP1 primary antibody (Cell Signaling, #17798) (**Supplemental Figure 9, Supplemental Table 1**), and fluorescence microscopy in the absence or presence of 200 µM sodium arsenite to induce SGs (**Supplemental Figure 9**).

### Creation of AAVS1 Safe Harbor LoxP Cassette Cells

The AAVS1 safe harbor donor plasmid containing the loxP-flanked blasticidin resistance cassette was a gift from the Taliaferro lab (University of Colorado). EGFP-G3BP1 ADAR1 KO and ADAR1p150 KO cells were split into a 48-well plate in 300 µL of media at a seeding density of 40,000 cells per well and allowed to incubate for 12-18 hours before transfection.

RNPs from Synthego with AAVS1 targeting sgRNA (**Supplemental Table 1**) were co-transfected into EGFP-G3BP1 ADAR1KO and ADAR1p150 KO cells using Lipofectamine LTX reagent along with the AAVS1 donor plasmid. The conditions were as follows: the first mixture contained 22 µL OPTI-MEM, 1.3 µL of 100 µM AAVS1 sgRNA, and 1 µL of Cas9, and 0.3 µg of the AAVS1 donor plasmid; the second contained 1.5 µL of LTX lipofectamine reagent and 22 µL of OPTI-MEM. The individual mixtures were vortexed and then allowed to incubate at room temperature for 10 minutes, before combining, vortexing, and allowed to incubate for another 10 minutes before adding to cells. A negative control with all components except the sgRNA was also carried out to ensure there were not off-target effects from Cas9 cleavage. The media was refreshed the following day, after which the cells were incubated for another 24 hr before splitting cells into fresh wells and beginning selection with blasticidin (6 μg mL^-1^ for the first two days followed by 10 μg mL^-1^ afterwards) until untransfected negative control cells had died and the newly selected loxP line exhibited normal growth patterns. Integration of the loxP cassette into the AAVS1 safe harbor was validate by PCR amplifying the junction of the AAVS1 site upstream of the inserted cassette. Primers (**Supplemental Table 1**) were designed such that products were only produced if the cassette was introduced as expected.

### Creation of ADAR1p150 Mutant Cell Lines Using Cre Recombination

The generation of the pRD-RIPE donor vector has been described elsewhere^87^. The Cre recombinase mammalian expression plasmid was acquired from addgene (pBT140, addgene #27493). The wild-type ADAR1p150-TST construct was synthesized and inserted into pRD-RIPE in place of the EGFG ORF by GenScript. All other mutant constructs were derived through mutagenesis of this plasmid, and were confirmed by full plasmid nanopore sequencing (Plasmidsaurus). The pRD-RIPE plasmid with doxycycline inducible ADAR1p150 transgenes also contained a constitutively active copy of the tetracycline transactivator and a puromycin resistance gene that replaces the blasticidin resistance gene in the AAVS1 safe harbor.

All loxP-containing cell lines followed the same protocol for cassette swapping. EGFP-G3BP1 ADAR1KO and ADAR1p150 KO cells were seeded into 6-well plates at a seeding density of 300,000 cells per well in a total of 2 mL of media and allowed to incubate for 12-18 hours before transfection. Cells were then co-transfected with 2 µg of pRD-RIPE donor plasmid (containing the different ADAR1p150 constructs) and 2% (wt/wt) of pBT140 (40 ng) using lipofectamine LTX. The same volumes of LTX components were used as for creation of EGFP-G3BP1 cell lines, see above. A negative control without the Cre-expressing plasmid was done to ensure that puromycin resistance was due to exchange into the AAVS1 safe harbor loxP cassette and not due to random integration. After 24 hours, the media was refreshed. Following another 24 hours, the cells were split into new wells and puromycin was added to 1 µg mL^-1^ to begin selection.

Incubation with puromycin proceeded for several days until control cells died and visible puromycin-resistant colonies began to emerge in the experimental wells. At this point, the concentration of puromycin was increased to 1.5 µg mL^-1^ and cells were expanded until complete recovery and new lines exhibited normal growth (**Supplemental Figure 1B**). Following selection, aliquots of newly created cell lines were frozen and stored.

### Dox Induction Optimization

A DOX concentration test (0, 1, 2, 5, 7.5, 10, 20, 50, 75, 100, 500, and 1000 ng mL^-1^ DOX) was carried out to optimize the expression level of the induced ADAR1p150 constructs to that of IFNβ (stem cell technologies) induced wild-type levels (**Figure 2B**). 50 ng mL^-1^ gave the closest expression level of ADAR1p150 transgenes compared to wild-type as judged by western blot (**Supplemental Figure 3**).

### Total RNA Extraction

The EGFP-G3BP1 AAVS1 loxP cassette cells expressing the different ADAR1p150 constructs were seeded into 6-well plates at a seeding density of 300,000 cells per well in 2 mL of media. Each cell line was processed in triplicate. 24 hours later, the cells were treated with 20 ng mL^-1^ IFNB (stem cell technologies) and either 50 ng mL^-1^ or 1 µg mL^-1^ DOX to induce ADAR1p150 expression and allowed to incubate for 24 hours before extraction. Cells were washed with 1 mL of ice cold 1X PBS to remove residual medium and detached cells and then RNA was extracted directly from the confluent 6-well plates using 500 µL of Trizol reagent (Invitrogen) and the samples were allowed to incubate for 5 min at room temperature. 100 μL of chloroform was added and the samples transferred to phase-lock tubes, centrifuged at 12,000 g for 15 min at 4°C, and the aqueous phase was transferred to a new tube. The aqueous phase was then combined with 300 μL of 70% EtOH, and transferred to an RNAeasy spin column, followed by centrifuging for 15 sec at 8000 g. The flow-through was discarded, and 350 µL of RW1 buffer was applied to the columns. The columns were spun for 15 sec at 8000, and the flow-through was again discarded. 10 µL of Qiagen DNase I was mixed with 70 µL of buffer RDD (per sample), applied to the column, and allowed to incubate for 30 min at room temperature to ensure digestion of genomic DNA. 350 µL of RW1 buffer was added to each column, allowed to incubate at room temperature for 5 min, and then spun down for 15 sec at 8000 g and the flow-through discarded. 500 µL of RPE was added to each column, and again centrifuged for 15 sec at 8000 g and the flow-through discarded. This was done twice for a total of two washes. The columns were placed into fresh collection tubes and centrifuged for 2 min at max speed to dry the column. The columns were then placed into 1.5 mL Eppendorf tubes, 30 µL of RNase-free water was applied directly to the membrane and allowed to incubate for 1 min at room temperature. The columns were then spun down at max speed for 1 min to elute the RNA. Purified RNA integrity and concentrations were verified using a NanoDrop Spectrophotometer (ThermoFisher), Qubit (ThermoFisher), and an electronic TapeStation 4150 System (Agilent). Samples with an RNA Integrity Number (RIN) ≥ 9 was selected for downstream processing.

### Western Blotting

At the same time as RNA-extraction, additional 6-well plates were seeded for western blotting of the expressed ADAR1p150 constructs, each sample in triplicate. Cells were washed with 1 mL of ice cold 1X PBS and then lysed in 400 µL of ice-cold RIPA lysis buffer containing 1X protease inhibitor cocktail and 1 mM phenylmethylsulphonyl fluoride (PMSF) (SantaCruz Biotechnology) directly in the plates. The plates were kept on ice for 10 minutes to allow complete lysis. The lysates were then transferred to microcentrifuge tubes and centrifuged at 14,000 g for 15 min to pellet cellular debris and the supernatant transferred to fresh microcentrifuge tubes. Protein concentrations were determined using a Pierce 660 nm Protein Assay (ThermoFisher) following the manufactures instructions. Protein concentrations were normalized to ensure equal loading across all samples.

Lysate samples were mixed with SDS loading buffer to 1X and then heated at 95°C for 5 minutes to ensure complete denaturation of proteins. Samples were then briefly centrifuged, and 15 µL of each sample was loaded onto a Mini-PROTRAN TGX gradient polyacrylamide (4-20%) gel (Bio-Rad) and electrophoresed at 180 volts in 1X SDS running buffer until the bromophenol blue dye front ran off the gel. Following electrophoresis, proteins were transferred onto a nitrocellulose membrane (MilliporeSigma) using a TransBlot semi-dry transfer system (Bio-Rad) at 10 V for 2 hours. The membrane was blocked for 1 hour at room temperature in 1X blocking buffer (PBS, 0.2% Tween (PBS-T), and 3% BSA). After blocking, the membrane was incubated overnight at 4°C with a rabbit anti-ADAR1 primary antibody (Cell Signaling, #14175) diluted 1:1000 in blocking buffer. Following incubation, the membrane was washed three times with 1X PBS-T (10 minutes each) and then incubated for 1 hour at room temperature with IRDye 800CW goat anti-rabbit secondary antibody (Licor-Bio) at a 1:10,000 dilution in blocking buffer, after which the membrane was washed three times with 1X PBS-T. The blot was imaged using a ChemiDoc MP (Bio-Rad) with detection set for the 800 nm channel. Afterwards, the membrane was incubated with mouse anti-β-actin antibody (Cell Signaling, #3700) overnight in blocking buffer, washed three times with PBS-T, incubated with goat IRDye 800CW goat anti-mouse secondary antibody (Licor-Bio) for 1 hour, washed three times again with PBS-T, and imaged using ChemiDoc MP. Western blots of ADAR1p150 were carried out identically as above using a rabbit anti-ADAR1p150 antibody (Cell Signaling, #32136).

Following imaging, protein band intensities were quantified using Fiji (ImageJ). The intensity values were normalized to β-actin, averaged over the three replicates, and the standard deviation was calculated to represent the variability between replicates. To determine the statistical significance between the expression levels of the integrated ADAR1p150 constructs and wild-type control cells, p-values were calculated using a two-tailed t-test. A p-value ≤ 0.05 (*), 0.01 (**), or 0.001 (***) was considered statistically significant.

### RNA-Seq Library Preparation

RNA-Seq libraries were prepared from 500 ng of total RNA extracted from each cell line using the Zymo-Seq RiboFree Total RNA Library Kit (Zymo Research), following the manufacturer’s protocol as provided with the version of the kit in use at the time. cDNA synthesis was performed from the total RNA, followed by RiboFree Universal Depletion for 1 hour at 98°C to enzymatically remove ribosomal RNA (rRNA), enriching for non-ribosomal RNA species.

Partial P5 and P7 adapters were then simultaneously ligated to the cDNA. Libraries were amplified by PCR for 12 cycles using unique dual indexing primers. Final libraries were purified using Select-a-Size MagBeads, quantified using a Qubit dsDNA High Sensitivity Assay, and assessed for fragment size distribution and absence of adapter dimers on an Agilent TapeStation. Libraries with appropriate concentration and size profiles were selected for sequencing.

### RNA-Sequencing

Sequencing was performed by Novogene. Paired-end sequencing (2x 150 bp) was conducted on a NovaSeq XPlus platform, generating at a read depth of ∼50 million reads per sample. All samples were pooled for multiplexing and subsequently sequenced on two independent flow cell lanes to ensure adequate coverage. Following sequencing, data from the two independent flow cells were retrieved, and corresponding reads were evaluated for potential lane-specific biases to ensure consistency across lanes before concatenating them into a single dataset per sample for downstream analysis.

### Mapping of RNA-seq Reads

Raw RNA-seq reads were first processed using Cutadapt v1.8.3^140^ to remove Illumina adaptor sequences and quality assessment was performed using FastQC v0.11.9. Reads were then mapped to the human reference genome (GRCh38) using STAR v2.7.10^141^ with default settings and the following additional parameters: “--alignSJoverhangMin 8 --alignSJDBoverhangMin 1 -- outFilterMismatchNoverLmax 0.04 --alignIntronMin 20 --alignIntronMax 1000000 -- alignMatesGapMax 1000000”. The STAR alignment was run in two-pass mode to first detect splice junctions and then realign reads for more accurate splicing event detection. The output files were generated in BAM format and unmapped reads were also collected for further analysis. We only considered uniquely mapped paired-end reads with a mapping quality of q=255. Duplicate alignments were marked with MarkDuplicates from Picard v2.7.0.

### Gene Expression Analysis

The RNA-seq data was processing using Salmon v1.9.0^89^ for transcript-level quantification with the Gencode v35 transcriptome assembly. Salmon was run in quasi-mapping-based mode using the index built from the Gencode v35 transcriptome and was executed with the following parameters: “--libType A, --validateMappings, and --numBootstraps 50”. Bootstrapping was performed 50 times to estimate the uncertainty of transcript abundance measurements. Gene expression levels were assessed via transcript per million (TPM) values and estimate counts for each transcript. Expression of the integrated ADAR1p150 constructs was carried out by including the sequences of the transgenes along with the reference transcriptome.

### Identification of A-to-I Editing Sites

Following alignment, BAM files produced by STAR were sorted and indexed using Samtools v1.16.1^142^. For detection of de novo A-to-I editing sites, we used the raer package (https://bioconductor.org/packages/3.19/bioc/html/raer.html) within an R-based pipeline. BAM files produced by STAR were sorted and indexed using Samtools v1.16.1135. BAM files were processed using pileup_sites from the raer package to generate a read count matrix for candidate editing sites. To ensure high confidence in editing site calls, we applied the following filtering parameters: minimum base quality of 30, minimum mapping quality of 255, and a minimum read depth of 5. Additional trimming of 5 nucleotides at the 5’ and 3’ ends was applied, and sites near indels (within 4 bp) were excluded. Spliced read alignments were required to have at least 10 bases aligned adjacent to the candidate editing site. Additionally, reads were excluded if they contained greater than 2 different types of variant base (e.g. A->G, G->T, C->A) or if the read base qualities were less than 20, in greater than 25% of the read. Variant detection was restricted to variants present in at least 2 reads and with an allelic frequency greater than 1%, and was quantified using at most 1 million reads. The RNA-seq libraries were stranded (fr-first-strand) and therefore the strandedness of the variant was assigned with sites from read 1 being anti-sense to the RNA strand, and sites from read 2 as sense to the RNA strand. Candidate sites with multiple alleles passing filters were excluded. To further focus on high-confidence A-to-I editing events, editing frequencies were computed for each site, and variants found in more than 3 samples with an editing frequency between 5% and 95% were kept for downstream analysis. We then excluded mitochondrial variants and those located within repetitive elements, such as simple repeats and low complexity regions, using repeat masking data from the UCSC genome browser (RepeatMasker track). Additionally, SNPs from dbSNP build 155 were annotated, and sites matching known SNPs were discarded. The remaining sites were considered as de novo A-to-I editing events.

REDIportal-detected sites were determined using the REDItoolDNARNA.py script provided as a part of the REDItools package v1.3^143^ using the following parameters: “-s 1 -q 10 -bq 20 -mbp 7 −l 20 -men 1 -T 1 -C -H”. Editing sites that were supported by fewer than 20 reads were filtered out. Putative editing sites were filtered against common SNPs from the dbSNP 155 to remove sequence variants. Editing events for sites present in the REDIportal database^143^ were enumerated using the raer package. Reads supporting an editing event or the reference sequence were counted for reads aligned with a minimum MAPQ of 255, and a minimum base quality of 30. Sites located in the first or last 5 bases of the read were excluded due to high alignment error rates in these read regions. The ALU editing index (AEI) was calculated using the raer function calc_AEI, based upon the algorithm developed by Roth et al. (REF, https://doi.org/10.1038/s41592-019-0610-9). ALU repeats were identified using the repeatMasker annotations provided by the UCSC genome browser. Known SNPs within ALU repeats were identified using the dbSNP 155 database, and excluded from the AEI calculation. Reads and base calls were filtering using the same criteria used for quantifying editing sites in REDIportal.

Hyper-edited regions were identified following a previously published pipeline and were implemented using custom Python scripts organized into a snakemake pipeline^144^. Unaligned reads from the STAR mapping were processed further by removing low-quality reads and those with repetitive sequences, following established protocols. After filtering, in forward-strand reads (from paired-end R2), adenines (A) were substituted with guanines (G) and aligned to a reference genome where adenines had similarly been converted to guanines to detect sense strand alignments, or to a genome where thymines (T) were converted to cytosines (C) for antisense detection. The reverse-strand reads (from R1) underwent a parallel process where thymines were changed to cytosines and aligned against genomes with either T-to-C or A-to-G conversions. Alignments were conducted using BWA v0.7.17 allowing up to two mismatches without introducing gaps^145^. Following alignment, mismatches between the original read sequences and the genome were evaluated. High-quality mismatches that met strict filtering criteria were classified as hyper-edited reads, as previously described^144^. This process was repeated for all 12 potential mismatch combinations to evaluate specificity for A-to-G changes. Hyper-edited segments within each read were defined as the interval from the first to the last mismatch, while hyper-edited clusters were identified by merging overlapping regions or those within 20 nucleotides. Clusters or sites supported by fewer than two reads were excluded. Editing frequencies were then calculated for hyper-edited sites not present in REDIportal, using STAR-aligned reads to the unmodified genome, with reference and alternative alleles counted with the raer package, using the same parameters as used for REDIportal sites.

The editing events identified from the REDIportal database were merged with those detected through the hyper-editing pipeline to generate a complete dataset representing all editing events. To identify editing clusters, we employed an approach recently employed by the Li lab^2^. We took the location information of editing sites in the genome as proxy for dsRNAs and applied a density-based clustering algorithm (DBSCAN) to identify genomic regions with many editing sites close together and editing indices across each cluster were calculated. For each cluster containing n editing sites, its editing index was calculated as follows: Σ_1 to n_ (G reads) / Σ_1 to n_ (A + G reads). To identify changes in editing frequencies between ADAR1p150 constructs, differential editing analysis was performed using DESeq2^104^. Editing sites were first filtered to include only those with sufficient sequencing depth (≥ 5 reads in all replicates within a comparison) and detectable editing activity (≥ 5% editing in at least three samples). For each editing site passing these filters, the number of edited reads and total reads were modeled as count data using a negative binomial distribution. Biological replicates (n = 3 per condition) were used to estimate within-group variability and to improve dispersion estimation. Pairwise comparisons between conditions were defined a priori, and for each comparison, a generalized linear model was fit to test whether editing frequencies differed significantly between the two conditions. Statistical significance was assessed using Wald tests, and resulting p-values were adjusted for multiple testing using the Benjamini-Hochberg procedure to control the false discovery rate. Editing sites with an adjusted p-value < 0.05 were considered significantly differentially edited.

For the plots of differentially-edited clusters, Alu foldbacks predicted by the CRSSANT software package^105^ were examined to identify foldbacks that were also ADAR1p150-dependent (ie, they have differential editing when comparing the ADAR1p150* plus and minus doxycycline conditions). Editing sites were only plotted on the predicted dsRNAs if they had an average editing frequency of > 0.05 averaged over the 3 replicates.

### Immunofluorescence and Microscopy

Cells were seeded into 24-well plates on 12 mm diameter, #1.5 thick polyD lysine-coated coverslips (NeuVitro) in media at a seeding density of 50,000 cells per cell (500 µL of media per well) and allowed to incubate for 24 hours. The cells were then treated with 20 ng mL^-1^ IFNB (stem cell technologies) and either 50 ng mL^-1^ or 1 µg mL^-1^ DOX to induce ADAR1p150 expression and allowed to incubate for another 24 hours. To induce SG formation, the media was aspirated and replaced with DMEM +FBS containing 200 µM sodium arsenite at hr 23 and allowed to incubate for 1 hr prior to the following steps. Cells were washed with 1 mL 1X PBS to remove residual medium and detached cells, and then fixed with 4% formaldehyde for 15 minutes at room temperature. Cells were then washed with 1X PBS and incubated with 1X PBS, 0.1% triton for 15 minutes at room temperature to permeabilize cells, after which they were incubated with blocking buffer (1X PBS, 5% BSA) for 1 hour at room temperature. After blocking, they were incubated with either rabbit anti-ADAR1 (Cell Signaling, #14175) or rabbit anti-ADAR1p150 (Cell Signaling, #32136) primary antibodies at a 1:100 dilution in overnight at 4°C. The cells were then washed three times with 1X PBS for 5 minutes each and then incubated with Alexa Fluor 594 conjugated goat anti-rabbit secondary antibody (Cell Signaling, #8889) at a dilution of 1:500 in blocking buffer for 1 hour at room temperature. The cells were washed three times with 1X PBS, before mounting on microscope slides using Vectashield mounting medium with DAPI (Vector Laboratories).

For the mCherry-tagged constructs, the cells were immediately fixed with 4% formaldehyde for 15 minutes, washed 3X with PBS, and then mounted on slides using Vectashield.

Z-stack images were taken on a DeltaVision Ultra with a 60x/1.42 (lens ID 10612) objective with 1.5200 oil immersion, and a charge coupled devide EDGE sCMOS camera. Cells were imaged over ∼50 focal planes of 0.2 μm. DAPI was imaged with a 10% transmittance and an exposure of 0.015 s, GFP was imaged with a 80% transmittance and an exposure of 0.010 s, and Alexa Fluor 594 was imaged with a 80% transmittance and an exposure of 0.025 s.

The cytoplasmic versus nuclear localization of the ADAR1p150 constructs was carried out by quantifying the Alexa Fluor 594 (red channel) intensity (through measuring the internal density) of the captured images using Fiji. The cytoplasmic and nuclear localization was calculated for the nucleus and cytoplasm for ∼30 cells per mutant and the average and standard deviation was reported. p-values were calculated using a two-sample homoscedastic two-tailed t-test. A p-value ≤ 0.05 (*), 0.01 (**), 0.001 (***), or 0.0001 (****) was considered statistically significant.

## DATA AND CODE AVAILABILITY

RNA-seq data will be deposited in the Gene Expression Omnibus and custom code used for RNA editing analyses will be made public under accession number GSE297017. The raer R package, used in this study for analyzing RNA editing patterns using RNA-seq data, is available on Bioconductor (https://bioconductor.org/packages/release/bioc/html/raer.html) and Github (https://github.com/rnabioco/raer).

## ASSOCIATED CONTENT

### Supporting Information

Supplemental Figures 1 through 11.

## AUTHOR INFORMATION

### Author Contributions

P.N. carried out all experiments and data analysis except for analysis of RNA-seq data, which was carried out by K.R.; J.K., M.H., J.R., and R.G. contributed intellectually to the project. P.N. wrote the manuscript with input from Q.V. and B.V.; all authors edited the manuscript; all authors have given approval to the final version of the manuscript.

## ACKNOWLEDGMENTS

We would like to thank the Rice lab (Rockefeller University) for generously sharing their ADAR1 and ADAR1p150 KO cell lines, as well as Na Zhao for feedback on the manuscript.

## Bibliography

1. Sadeq, S., Al-Hashimi, S., Cusack, C. M. & Werner, A. Endogenous double-stranded RNA. Noncoding RNA (2021) doi:10.3390/ncrna7010015.

2. Sun, T. et al. A Small Subset of Cytosolic dsRNAs Must Be Edited by ADAR1 to Evade MDA5-Mediated Autoimmunity. bioRxiv 2022.08.29.505707 (2022) doi:10.1101/2022.08.29.505707.

3. Chen, Y. G. & Hur, S. Cellular origins of dsRNA, their recognition and consequences. Nat Rev Mol Cell Biol 23, 286–301 (2022).

4. Medstrand, P., Van De Lagemaat, L. N. & Mager, D. L. Retroelement distributions in the human genome: Variations associated with age and proximity to genes. Genome Res (2002) doi:10.1101/gr.388902.

5. Deininger, P. Alu elements: Know the SINEs. Genome Biology Preprint at 10.1186/gb-2011-12-12-236 (2011).

6. Salem, A. H. et al. Alu elements and hominid phylogenetics. Proc Natl Acad Sci U S A (2003) doi:10.1073/pnas.2133766100.

7. Rehwinkel, J. & Gack, M. U. RIG-I-like receptors: their regulation and roles in RNA sensing. Nature Reviews Immunology Preprint at 10.1038/s41577-020-0288-3 (2020).

8. Goubau, D., Deddouche, S. & Reis e Sousa, C. Cytosolic Sensing of Viruses. Immunity Preprint at 10.1016/j.immuni.2013.05.007 (2013).

9. Gong, C. & Maquat, L. E. LncRNAs transactivate STAU1-mediated mRNA decay by duplexing with 39 UTRs via Alu eleme. Nature (2011) doi:10.1038/nature09701.

10. Ahmad, S. et al. Breaching Self-Tolerance to Alu Duplex RNA Underlies MDA5-Mediated Inflammation. Cell (2018) doi:10.1016/j.cell.2017.12.016.

11. Mehdipour, P. et al. Epigenetic therapy induces transcription of inverted SINEs and ADAR1 dependency. Nature (2020) doi:10.1038/s41586-020-2844-1.

12. Aloni, Y. & Attardi, G. Symmetrical in vivo transcription of mitochondrial DNA in HeLa cells. Proc Natl Acad Sci U S A (1971) doi:10.1073/pnas.68.8.1757.

13. Young, P. G. & Attardi, G. Characterization of double-stranded RNA from HeLa cell mitochondria. Biochem Biophys Res Commun (1975) doi:10.1016/S0006-291X(75)80357-3.

14. D’Souza, A. R. & Minczuk, M. Mitochondrial transcription and translation: Overview. Essays in Biochemistry Preprint at 10.1042/EBC20170102 (2018).

15. Werner, A., Kanhere, A., Wahlestedt, C. & Mattick, J. S. Natural antisense transcripts as versatile regulators of gene expression. Nat Rev Genet (2024) doi:10.1038/s41576-024-00723-z.

16. Ali, A. et al. Therapeutic potential of natural antisense transcripts and various mechanisms involved for clinical applications and disease prevention. RNA Biology Preprint at 10.1080/15476286.2023.2293335 (2023).

17. Böttcher, R., Schmidts, I., Nitschko, V., Duric, P. & Förstemann, K. RNA polymerase II is recruited to DNA double-strand breaks for dilncRNA transcription in Drosophila. RNA Biol (2022) doi:10.1080/15476286.2021.2014694.

18. López-Polo, V. et al. Release of mitochondrial dsRNA into the cytosol is a key driver of the inflammatory phenotype of senescent cells. Nat Commun 15, 7378 (2024).

19. Yu, Q., Herrero del Valle, A., Singh, R. & Modis, Y. MDA5 disease variant M854K prevents ATP-dependent structural discrimination of viral and cellular RNA. Nat Commun (2021) doi:10.1038/s41467-021-27062-5.

20. Kato, H. et al. Length-dependent recognition of double-stranded ribonucleic acids by retinoic acid-inducible gene-I and melanoma differentiation-associated gene 5. Journal of Experimental Medicine (2008) doi:10.1084/jem.20080091.

21. Peisley, A. et al. Kinetic mechanism for viral dsRNA length discrimination by MDA5 filaments. Proc Natl Acad Sci U S A (2012) doi:10.1073/pnas.1208618109.

22. Yu, Q., Qu, K. & Modis, Y. Cryo-EM Structures of MDA5-dsRNA Filaments at Different Stages of ATP Hydrolysis. Mol Cell (2018) doi:10.1016/j.molcel.2018.10.012.

23. Bass, B. L. RNA editing by adenosine deaminases that act on RNA. Annual Review of Biochemistry Preprint at 10.1146/annurev.biochem.71.110601.135501 (2002).

24. Chung, H. et al. Human ADAR1 Prevents Endogenous RNA from Triggering Translational Shutdown. Cell (2018) doi:10.1016/j.cell.2017.12.038.

25. Liddicoat, B. J. et al. RNA editing by ADAR1 prevents MDA5 sensing of endogenous dsRNA as nonself. Science (1979) (2015) doi:10.1126/science.aac7049.

26. Rodero, M. P. & Crow, Y. J. Type I interferon–mediated monogenic autoinflammation: The type I interferonopathies, a conceptual overview. Journal of Experimental Medicine (2016) doi:10.1084/jem.20161596.

27. Dias Junior, A. G., Sampaio, N. G. & Rehwinkel, J. A Balancing Act: MDA5 in Antiviral Immunity and Autoinflammation. Trends in Microbiology Preprint at 10.1016/j.tim.2018.08.007 (2019).

28. Hartner, J. C. et al. Liver Disintegration in the Mouse Embryo Caused by Deficiency in the RNA-editing Enzyme ADAR1. Journal of Biological Chemistry (2004) doi:10.1074/jbc.M311347200.

29. Hartner, J. C., Walkley, C. R., Lu, J. & Orkin, S. H. ADAR1 is essential for the maintenance of hematopoiesis and suppression of interferon signaling. Nat Immunol (2009) doi:10.1038/ni.1680.

30. Wang, Q., Khillan, J., Gadue, P. & Nishikura, K. Requirement of the RNA editing deaminase ADAR1 gene for embryonic erythropoiesis. Science (1979) (2000) doi:10.1126/science.290.5497.1765.

31. Wang, Q. et al. Stress-induced Apoptosis Associated with Null Mutation of ADAR1 RNA Editing Deaminase Gene. Journal of Biological Chemistry (2004) doi:10.1074/jbc.M310162200.

32. Ward, S. V. et al. RNA editing enzyme adenosine deaminase is a restriction factor for controlling measles virus replication that also is required for embryogenesis. Proc Natl Acad Sci U S A (2011) doi:10.1073/pnas.1017241108.

33. Bass, B. L. & Weintraub, H. An unwinding activity that covalently modifies its double-stranded RNA substrate. Cell (1988) doi:10.1016/0092-8674(88)90253-X.

34. Wagner, R. W., Smith, J. E., Cooperman, B. S. & Nishikura, K. A double-stranded RNA unwinding activity introduces structural alterations by means of adenosine to inosine conversions in mammalian cells and Xenopus eggs. Proc Natl Acad Sci U S A (1989) doi:10.1073/pnas.86.8.2647.

35. Wright, D. J., Rice, J. L., Yanker, D. M. & Znosko, B. M. Nearest neighbor parameters for inosine x uridine pairs in RNA duplexes. Biochemistry 46, 4625–4634 (2007).

36. Serra, M. J., Smolter, P. E. & Westhof, E. Pronounced instability of tandem IU base pairs in RNA. Nucleic Acids Res 32, 1824–1828 (2004).

37. Banerjee, A., Anand, M., Kalita, S. & Ganji, M. Single-molecule analysis of DNA base-stacking energetics using patterned DNA nanostructures. Nat Nanotechnol (2023) doi:10.1038/s41565-023-01485-1.

38. Chung, H. et al. Human ADAR1 Prevents Endogenous RNA from Triggering Translational Shutdown. Cell (2018) doi:10.1016/j.cell.2017.12.038.

39. Athanasiadis, A., Rich, A. & Maas, S. Widespread A-to-I RNA editing of Alu-containing mRNAs in the human transcriptome. PLoS Biol (2004) doi:10.1371/journal.pbio.0020391.

40. Bazak, L. et al. A-to-I RNA editing occurs at over a hundred million genomic sites, located in a majority of human genes. Genome Res (2014) doi:10.1101/gr.164749.113.

41. Kim, D. D. Y. et al. Widespread RNA editing of embedded Alu elements in the human transcriptome. Genome Res (2004) doi:10.1101/gr.2855504.

42. Levanon, E. Y. et al. Systematic identification of abundant A-to-I editing sites in the human transcriptome. Nat Biotechnol 22, 1001–1005 (2004).

43. Porath, H. T., Carmi, S. & Levanon, E. Y. A genome-wide map of hyper-edited RNA reveals numerous new sites. Nat Commun (2014) doi:10.1038/ncomms5726.

44. Ramaswami, G. et al. Accurate identification of human Alu and non-Alu RNA editing sites. Nat Methods (2012) doi:10.1038/nmeth.1982.

45. Lander, E. S. et al. Initial sequencing and analysis of the human genome. Nature (2001) doi:10.1038/35057062.

46. Dhir, A. et al. Mitochondrial double-stranded RNA triggers antiviral signalling in humans. Nature (2018) doi:10.1038/s41586-018-0363-0.

47. Faghihi, M. A. & Wahlestedt, C. Regulatory roles of natural antisense transcripts. Nature Reviews Molecular Cell Biology Preprint at 10.1038/nrm2738 (2009).

48. Patterson, J. B., Thomis, D. C., Hans, S. L. & Samuel, C. E. Mechanism of interferon action: double-stranded RNA-specific adenosine deaminase from human cells is inducible by alpha and gamma interferons. Virology 210, 508–511 (1995).

49. Patterson, J. B. & Samual, S. E. Expression and regulation by interferon of a double-stranded-RNA-specific adenosine deaminase from human cells: evidence for two forms of the deaminase. Mol Cell Biol 15, 5376–5388 (1995).

50. George, C. X. & Samuel, C. E. Characterization of the 5’-flanking region of the human RNA-specific adenosine deaminase ADAR1 gene and identification of an interferon-inducible ADAR1 promoter. Gene (1999) doi:10.1016/S0378-1119(99)00017-7.

51. Strehblow, A., Hallegger, M. & Jantsch, M. F. Nucleocytoplasmic distribution of human RNA-editing enzyme ADAR1 is modulated by double-stranded RNA-binding domains, a leucine-rich export signal, and a putative dimerization domain. Mol Biol Cell 13, 3822– 3835 (2002).

52. George, C. X. & Samuel, C. E. Human RNA-specific adenosine deaminase ADAR1 transcripts possess alternative exon 1 structures that initiate from different promoters, one constitutively active and the other interferon inducible. Proc Natl Acad Sci U S A (1999) doi:10.1073/pnas.96.8.4621.

53. Eckmann, C. R., Neunteufl, A., Pfaffstetter, L. & Jantsch, M. F. The human but not the Xenopus RNA-editing enzyme ADAR1 has an atypical nuclear localization signal and displays the characteristics of a shuttling protein. Mol Biol Cell 12, 1911–1924 (2001).

54. Rice, G. I. et al. Mutations in ADAR1 cause Aicardi-Goutières syndrome associated with a type i interferon signature. Nat Genet (2012) doi:10.1038/ng.2414.

55. Pestal, K. et al. Isoforms of RNA-Editing Enzyme ADAR1 Independently Control Nucleic Acid Sensor MDA5-Driven Autoimmunity and Multi-organ Development. Immunity (2015) doi:10.1016/j.immuni.2015.11.001.

56. Kim, J. I. et al. RNA editing at a limited number of sites is sufficient to prevent MDA5 activation in the mouse brain. PLoS Genet (2021) doi:10.1371/journal.pgen.1009516.

57. Hall, K., Cruz, P., Tinoco, I., Jovin, T. M. & Van De Sande, J. H. ‘Z-RNA’ - A left-handed RNA double helix. Nature (1984) doi:10.1038/311584a0.

58. Schwartz, T., Rould, M. A., Lowenhaupt, K., Herbert, A. & Rich, A. Crystal structure of the Zα domain of the human editing enzyme ADAR1 bound to left-handed Z-DNA. Science (1979) (1999) doi:10.1126/science.284.5421.1841.

59. Placido, D., Brown 2nd, B. A., Lowenhaupt, K., Rich, A. & Athanasiadis, A. A left-handed RNA double helix bound by the Z alpha domain of the RNA-editing enzyme ADAR1. Structure 15, 395–404 (2007).

60. Popenda, M., Milecki, J. & Adamiak, R. W. High salt solution structure of a left-handed RNA double helix. Nucleic Acids Res 32, 4044–4054 (2004).

61. Wang, A. H. J. et al. Molecular structure of a left-Handed double helical DNA fragment at atomic resolution. Nature (1979) doi:10.1038/282680a0.

62. Ho, P. S. & Mooers, B. H. Z-DNA crystallography. Biopolymers 14, 65–90 (1997).

63. Wang, A. J. et al. Left-handed double helical DNA: variations in the backbone conformation. Science (1979) 211, 171–176 (1981).

64. Krall, J. B., Nichols, P. J., Henen, M. A., Vicens, Q. & Vögeli, B. Structure and Formation of Z-DNA and Z-RNA. Molecules 28, (2023).

65. Šponer, J., Gabb, H. A., Leszczynski, J. & Hobza, P. Base-base and deoxyribose-base stacking interactions in B-DNA and Z-DNA: A quantum-chemical study. Biophys J (1997) doi:10.1016/S0006-3495(97)78049-4.

66. Kollman, P. A., Weiner, P. K. & Dearing, A. THEORETICAL STUDIES OF THE STRUCTURE AND ENERGIES OF BASE-PAIRED NUCLEOTIDES AND THE DISSOCIATION KINETICS OF A PROFLAVINE-DINUCLEOTIDE COMPLEX. Ann N Y Acad Sci (1981) doi:10.1111/j.1749-6632.1981.tb50572.x.

67. Gueron, M. & Demaret, J. P. A simple explanation of the electrostatics of the B-to-Z transition of DNA. Proc Natl Acad Sci U S A (1992) doi:10.1073/pnas.89.13.5740.

68. Misra, V. K. & Honig, B. The electrostatic contribution to the B to Z transition of DNA. Biochemistry (1996) doi:10.1021/bi951463y.

69. Guéron, M., Demaret, J. P. & Filoche, M. A unified theory of the B-Z transition of DNA in high and low concentrations of multivalent ions. Biophys J (2000) doi:10.1016/S0006-3495(00)76665-3.

70. Brown 2nd, B. A., Lowenhaupt, K., Wilbert, C. M., Hanlon, E. B. & Rich, A. The zalpha domain of the editing enzyme dsRNA adenosine deaminase binds left-handed Z-RNA as well as Z-DNA. Proc Natl Acad Sci U S A 97, 13532–13536 (2000).

71. Schwartz, T., Rould, M. A., Lowenhaupt, K., Herbert, A. & Rich, A. Crystal structure of the Zalpha domain of the human editing enzyme ADAR1 bound to left-handed Z-DNA. Science (1979) 11, 1841–1845 (1999).

72. Nakahama, T. et al. Mutations in the adenosine deaminase ADAR1 that prevent endogenous Z-RNA binding induce Aicardi-Goutieres-syndrome-like encephalopathy. Immunity 54, 1976–1988 e7 (2021).

73. Tang, Q. et al. Adenosine-to-inosine editing of endogenous Z-form RNA by the deaminase ADAR1 prevents spontaneous MAVS-dependent type I interferon responses. Immunity 54, 1961–1975 (2021).

74. de Reuver, R. et al. ADAR1 interaction with Z-RNA promotes editing of endogenous double-stranded RNA and prevents MDA5-dependent immune activation. Cell Rep (2021) doi:10.1016/j.celrep.2021.109500.

75. Mannion, N. M. et al. The RNA-Editing Enzyme ADAR1 Controls Innate Immune Responses to RNA. Cell Rep (2014) doi:10.1016/j.celrep.2014.10.041.

76. Koeris, M., Funke, L., Shrestha, J., Rich, A. & Maas, S. Modulation of ADAR1 editing activity by Z-RNA in vitro. Nucleic Acids Res (2005) doi:10.1093/nar/gki849.

77. Xing, Y. et al. RNA editing of AZIN1 coding sites is catalyzed by ADAR1 p150 after splicing. Journal of Biological Chemistry (2023) doi:10.1016/j.jbc.2023.104840.

78. Kleinova, R. et al. The ADAR1 editome reveals drivers of editing-specificity for ADAR1-isoforms. Nucleic Acids Res (2023) doi:10.1093/nar/gkad265.

79. Guo, X. et al. ADAR1 Zα domain P195A mutation activates the MDA5-dependent RNA-sensing signaling pathway in brain without decreasing overall RNA editing. Cell Rep (2023) doi:10.1016/j.celrep.2023.112733.

80. Livingston, J. H. et al. A type I interferon signature identifies bilateral striatal necrosis due to mutations in ADAR1. J Med Genet 51, 76–82 (2014).

81. Herbert, A. Mendelian disease caused by variants affecting recognition of Z-DNA and Z-RNA by the Zα domain of the double-stranded RNA editing enzyme ADAR. European Journal of Human Genetics 28, 114–117 (2020).

82. Langeberg, C. J., Nichols, P. J., Henen, M. A., Vicens, Q. & Vögeli, B. Differential Structural Features of Two Mutant ADAR1p150 Zα Domains Associated with Aicardi-Goutières Syndrome. J Mol Biol 435, 168040 (2023).

83. Feng, S. et al. Alternate rRNA secondary structures as regulators of translation. Nat Struct Mol Biol (2011) doi:10.1038/nsmb.1962.

84. Tang, Q. et al. Adenosine-to-inosine editing of endogenous Z-form RNA by the deaminase ADAR1 prevents spontaneous MAVS-dependent type I interferon responses. Immunity 54, 1961–1975 (2021).

85. de Reuver, R. et al. ADAR1 interaction with Z-RNA promotes editing of endogenous double-stranded RNA and prevents MDA5-dependent immune activation. Cell Rep (2021) doi:10.1016/j.celrep.2021.109500.

86. Liu, Y. & Samuel, C. E. Mechanism of interferon action: functionally distinct RNA-binding and catalytic domains in the interferon-inducible, double-stranded RNA-specific adenosine deaminase. J Virol (1996) doi:10.1128/jvi.70.3.1961-1968.1996.

87. Khandelia, P., Yap, K. & Makeyev, E. V. Streamlined platform for short hairpin RNA interference and transgenesis in cultured mammalian cells. Proc Natl Acad Sci U S A (2011) doi:10.1073/pnas.1103532108.

88. Chung, H. et al. Human ADAR1 Prevents Endogenous RNA from Triggering Translational Shutdown. Cell (2018) doi:10.1016/j.cell.2017.12.038.

89. Patro, R., Duggal, G., Love, M. I., Irizarry, R. A. & Kingsford, C. Salmon provides fast and bias-aware quantification of transcript expression. Nat Methods 14, 417–419 (2017).

90. Gan, J. et al. USP16 is an ISG15 cross-reactive deubiquitinase that targets pro-ISG15 and ISGylated proteins involved in metabolism. Proc Natl Acad Sci U S A 120, (2023).

91. Guo, X. et al. An AGS-associated mutation in ADAR1 catalytic domain results in early-onset and MDA5-dependent encephalopathy with IFN pathway activation in the brain. J Neuroinflammation 19, 1–16 (2022).

92. Patterson, J. B. & Samual, S. E. Expression and regulation by interferon of a double-stranded-RNA-specific adenosine deaminase from human cells: evidence for two forms of the deaminase. Mol Cell Biol 15, 5376–5388 (1995).

93. Weier, H. U. G., George, C. X., Greulich, K. M. & Samuel, C. E. The Interferon-Inducible, Double-Stranded RNA-Specific Adenosine Deaminase Gene (DSRAD) Maps to Human Chromosome 1q21.1–21.2. Genomics 30, 372–375 (1995).

94. George, C. X. & Samuel, C. E. Human RNA-specific adenosine deaminase ADAR1 transcripts possess alternative exon 1 structures that initiate from different promoters, one constitutively active and the other interferon inducible. Proc Natl Acad Sci U S A (1999) doi:10.1073/pnas.96.8.4621.

95. George, C. X. & Samuel, C. E. Characterization of the 5′-flanking region of the human RNA-specific adenosine deaminase ADAR1 gene and identification of an interferon-inducible ADAR1 promoter. Gene 229, 203–213 (1999).

96. Mendoza, H. G. & Beal, P. A. Structural and functional effects of inosine modification in mRNA. RNA 30, 512–520 (2024).

97. Mansi, L. et al. REDIportal: Millions of novel A-to-I RNA editing events from thousands of RNAseq experiments. Nucleic Acids Res (2021) doi:10.1093/nar/gkaa916.

98. Blow, M., Futreal, A. P., Wooster, R. & Stratton, M. R. A survey of RNA editing in human brain. Genome Res (2004) doi:10.1101/gr.2951204.

99. Carmi, S., Borukhov, I. & Levanon, E. Y. Identification of widespread ultra-edited human RNAs. PLoS Genet (2011) doi:10.1371/journal.pgen.1002317.

100. Barak, M. et al. Evidence for large diversity in the human transcriptome created by Alu RNA editing. Nucleic Acids Res (2009) doi:10.1093/nar/gkp729.

101. Neeman, Y., Levanon, E. Y., Jantsch, M. F. & Eisenberg, E. RNA editing level in the mouse is determined by the genomic repeat repertoire. RNA (2006) doi:10.1261/rna.165106.

102. Nichols, P. J. et al. Z-Form Adoption of Nucleic Acid is a Multi-Step Process Which Proceeds through a Melted Intermediate. J Am Chem Soc (2023) doi:10.1021/jacs.3c10406.

103. Herzog, V. A. et al. Thiol-linked alkylation of RNA to assess expression dynamics. Nature Methods 2017 14:12 14, 1198–1204 (2017).

104. Love, M. I., Huber, W. & Anders, S. Moderated estimation of fold change and dispersion for RNA-seq data with DESeq2. Genome Biol 15, 550 (2014).

105. Zhang, M. et al. Classification and clustering of RNA crosslink-ligation data reveal complex structures and homodimers. Genome Res (2022) doi:10.1101/gr.275979.121.

106. Ng, S. K., Weissbach, R., Ronson, G. E. & Scadden, A. D. J. Proteins that contain a functional Z-DNA-binding domain localize to cytoplasmic stress granules. Nucleic Acids Res (2013) doi:10.1093/nar/gkt750.

107. Gabriel, L. et al. Enrichment of Zα domains at cytoplasmic stress granules is due to their innate ability to bind to nucleic acids. J Cell Sci (2021) doi:10.1242/jcs.258446.

108. Kedersha, N. et al. G3BP-Caprin1-USP10 complexes mediate stress granule condensation and associate with 40S subunits. Journal of Cell Biology (2016) doi:10.1083/jcb.201508028.

109. John, L. & Samuel, C. E. Induction of stress granules by interferon and down-regulation by the cellular RNA adenosine deaminase ADAR1. Virology (2014) doi:10.1016/j.virol.2014.02.025.

110. Poulsen, H., Nilsson, J., Damgaard, C. K., Egebjerg, J. & Kjems, J. CRM1 mediates the export of ADAR1 through a nuclear export signal within the Z-DNA binding domain. Mol Cell Biol 21, 7862–7871 (2001).

111. Roth, S. H., Levanon, E. Y. & Eisenberg, E. Genome-wide quantification of ADAR adenosine-to-inosine RNA editing activity. Nat Methods (2019) doi:10.1038/s41592-019-0610-9.

112. Nichols, P. J. et al. Zα Domain of ADAR1 Binds to an A-Form-like Nucleic Acid Duplex with Low Micromolar Affinity. Biochemistry (2024) doi:10.1021/acs.biochem.3c00636.

113. de Reuver, R. et al. ADAR1 prevents autoinflammation by suppressing spontaneous ZBP1 activation. Nature 631, (2022).

114. Yang, T. et al. Triggering endogenous Z-RNA sensing for anti-tumor therapy through ZBP1-dependent necroptosis. Cell Rep 42, 113377 (2023).

115. Wang, J., Wang, S., Zhong, C., Tian, T. & Zhou, X. Novel insights into a major DNA oxidative lesion: Its effects on Z-DNA stabilization. Org Biomol Chem (2015) doi:10.1039/c5ob01340b.

116. Vongsutilers, V. & Gannett, P. M. C8-Guanine modifications: Effect on Z-DNA formation and its role in cancer. Organic and Biomolecular Chemistry Preprint at 10.1039/c8ob00030a (2018).

117. Xu, Y., Ikeda, R. & Sugiyama, H. 8-Methylguanosine: a powerful Z-DNA stabilizer. J Am Chem Soc 125, 13519–13524 (2003).

118. Hahm, J. Y., Park, J., Jang, E. S. & Chi, S. W. 8-Oxoguanine: from oxidative damage to epigenetic and epitranscriptional modification. Experimental and Molecular Medicine Preprint at 10.1038/s12276-022-00822-z (2022).

119. Maelfait, J. & Rehwinkel, J. The Z-nucleic acid sensor ZBP1 in health and disease. Journal of Experimental Medicine Preprint at 10.1084/JEM.20221156 (2023).

120. Kuriakose, T. & Kanneganti, T. D. ZBP1: Innate Sensor Regulating Cell Death and Inflammation. Trends in Immunology Preprint at 10.1016/j.it.2017.11.002 (2018).

121. Bajad, P. et al. An internal deletion of ADAR rescued by MAVS deficiency leads to a minute phenotype. Nucleic Acids Res (2020) doi:10.1093/nar/gkaa025.

122. Hubbard, N. et al. ADAR1 mutation causes ZBP1-dependent immunopathology. Nature 607, 769–775 (2022).

123. Jiao, H. et al. ADAR1 averts fatal type I interferon induction by ZBP1. Nature (2022) doi:10.1038/s41586-022-04878-9.

124. Zhang, T. et al. ADAR1 masks the cancer immunotherapeutic promise of ZBP1-driven necroptosis. Nature 606, 594–602 (2022).

125. Karki, R. et al. ADAR1 restricts ZBP1-mediated immune response and PANoptosis to promote tumorigenesis. Cell Rep (2021) doi:10.1016/j.celrep.2021.109858.

126. Sijen, T. & Plasterk, R. H. A. Transposon silencing in the Caenorhabditis elegans germ line by natural RNAi. Nature (2003) doi:10.1038/nature02107.

127. Reich, D. P. & Bass, B. L. Mapping the dsrna world. Cold Spring Harb Perspect Biol (2019) doi:10.1101/cshperspect.a035352.

128. Herbert, A. et al. A Z-DNA binding domain present in the human editing enzyme, double-stranded RNA adenosine deaminase. Proc Natl Acad Sci U S A (1997) doi:10.1073/pnas.94.16.8421.

129. Nichols, P. J. et al. Zα Domain of ADAR1 Binds to an A-Form-like Nucleic Acid Duplex with Low Micromolar Affinity. Biochemistry (2024) doi:10.1021/acs.biochem.3c00636.

130. Nichols, P. J. et al. Z-Form Adoption of Nucleic Acid is a Multi-Step Process Which Proceeds through a Melted Intermediate. J Am Chem Soc (2023) doi:10.1021/jacs.3c10406.

131. Yi-Brunozzi, H. Y., Stephens, O. M. & Beal, P. A. Conformational Changes That Occur during an RNA-editing Adenosine Deamination Reaction. Journal of Biological Chemistry 276, 37827–37833 (2001).

132. Karki, A., Campbell, K. B., Mozumder, S., Fisher, A. J. & Beal, P. A. Impact of Disease-Associated Mutations on the Deaminase Activity of ADAR1. Biochemistry 63, 282–293 (2024).

133. Yang, W. et al. ADAR1 RNA Deaminase Limits Short Interfering RNA Efficacy in Mammalian Cells. J Biol Chem 280, 3946 (2004).

134. Nichols, P. J. et al. Recognition of non-CpG repeats in Alu and ribosomal RNAs by the Z-RNA binding domain of ADAR1 induces A-Z junctions. Nat Commun (2021) doi:10.1038/s41467-021-21039-0.

135. Schwartz, T., Rould, M. A., Lowenhaupt, K., Herbert, A. & Rich, A. Crystal structure of the Zα domain of the human editing enzyme ADAR1 bound to left-handed Z-DNA. Science (1979) (1999) doi:10.1126/science.284.5421.1841.

136. Placido, D., Brown, B. A., Lowenhaupt, K., Rich, A. & Athanasiadis, A. A Left-Handed RNA Double Helix Bound by the Zα Domain of the RNA-Editing Enzyme ADAR1. Structure (2007) doi:10.1016/j.str.2007.03.001.

137. Romero, M. F. et al. Novel Z-DNA binding domains in giant viruses. J Biol Chem 300, 107504 (2024).

138. Labun, K. et al. CHOPCHOP v3: Expanding the CRISPR web toolbox beyond genome editing. Nucleic Acids Res (2019) doi:10.1093/nar/gkz365.

139. Cong, L. et al. Multiplex genome engineering using CRISPR/Cas systems. Science (1979) (2013) doi:10.1126/science.1231143.

140. Martin, M. Cutadapt removes adapter sequences from high-throughput sequencing reads. EMBnet J (2011) doi:10.14806/ej.17.1.200.

141. Dobin, A. et al. STAR: Ultrafast universal RNA-seq aligner. Bioinformatics (2013) doi:10.1093/bioinformatics/bts635.

142. Danecek, P. et al. Twelve years of SAMtools and BCFtools. Gigascience (2021) doi:10.1093/gigascience/giab008.

143. Picardi, E. & Pesole, G. REDItools: High-throughput RNA editing detection made easy. Bioinformatics (2013) doi:10.1093/bioinformatics/btt287.

144. Pinto, Y., Cohen, H. Y. & Levanon, E. Y. Mammalian conserved ADAR targets comprise only a small fragment of the human editosome. Genome Biol (2014) doi:10.1186/gb-2014-15-1-r5.

145. Li, H. & Durbin, R. Fast and accurate short read alignment with Burrows-Wheeler transform. Bioinformatics (2009) doi:10.1093/bioinformatics/btp324.

